# Testing the ability of species distribution models to infer variable importance

**DOI:** 10.1101/715904

**Authors:** Adam B. Smith, Maria J. Santos

## Abstract

Models of species’ distributions and niches are frequently used to infer the importance of range- and niche-defining variables. However, the degree to which these models can reliably identify important variables and quantify their influence remains unknown. Here we use a series of simulations to explore how well models can 1) discriminate between variables with different influence and 2) calibrate the magnitude of influence relative to an “omniscient” model. To quantify variable importance, we trained generalized additive models (GAMs), Maxent, and boosted regression trees (BRTs) on simulated data and tested their sensitivity to permutations in each predictor. Importance was inferred by calculating the correlation between permuted and unpermuted predictions, and by comparing predictive accuracy of permuted and unpermuted predictions using AUC and the Continuous Boyce Index. In scenarios with one influential and one uninfluential variable, models were unable to discriminate reliably between variables in conditions that are normally challenging for generating accurate predictions: training occurrences <8-64; prevalence >0.5; small spatial extent; environmental data with coarse resolution when spatial autocorrelation is low; and correlation between environmental variables where |*r*| >0.7. When two variables influenced the distribution equally, importance was underestimated when species had narrow or intermediate niche breadth. Interactions between variables in how they shaped the niche did not affect inferences about their importance. When variables acted unequally, the effect of the stronger variable was overestimated. GAMs and Maxent discriminated between variables more reliably than BRTs, but no algorithm was consistently well-calibrated vis-à-vis the omniscient model. Algorithm-specific measures of importance like Maxent’s change-in-gain metric were less robust than the permutation test. Overall, high predictive accuracy did not connote robust inferential capacity. As a result, requirements for reliably measuring variable importance are likely more stringent than for creating models with high predictive accuracy.

## Introduction

What environmental factors determine species’ ranges and environmental tolerances? The answer to this question remains elusive for most species on Earth despite being crucial for addressing long-standing issues in theoretical and applied ecology. Although manipulative field- and laboratory-based studies can identify factors that shape niches and ranges (Hargreaves et al. 2016; Lee-Yaw et al. 2016), for most species logistical difficulties preclude examining large numbers of variables and studying range limits across wide geographic scales. Alternatively, environmental limits can be inferred using models of species’ geographic ranges and niches. These ecological niche models and species distribution models (SDMs) are constructed by correlating observations of occurrence with data on environmental conditions at occupied sites. Indeed, one of the most common uses for SDMs is to identify important variables (e.g., 227 inferential studies analyzed by Bradie & Leung 2017 using the Maxent algorithm alone). However, we are aware of no studies that systematically evaluate how well SDMs measure variable importance. Compared to the attention devoted to understanding predictive accuracy of models of niches and distributions (e.g., Elith et al. 2006; Smith et al. 2013; Buklin et al. 2015; Guevara et al. 2018; Norberg et al. 2019), the lack of research on the efficacy of these same models for identifying important variables is a striking oversight.

The ability of an SDM to infer variable importance will likely be affected by factors that are extrinsic to the species and thus at least nominally under control of the modeler (e.g., sample size, study region extent, etc.) and by factors intrinsic to the species (e.g., niche breadth). Here we utilize a reductionist approach based on virtual species (Meynard et al. 2019) to systematically evaluate a set of extrinsic and intrinsic factors expected to influence variable inference. Our goal was to identify the minimal conditions under which each factor allows robust inference of variable importance, assuming all other conditions are optimal. We conducted nine simulation experiments to explore circumstances that affect inference. We start with the simplest case in which a species’ range is determined by a single “TRUE” variable, but the SDMs are presented with data on this variable plus an uncorrelated “FALSE” variable with no effect on distribution. We then explore the effects of sample size, spatial scale, and collinearity between variables. Finally, we examine cases where the species’ distribution is shaped by two collinear TRUE variables that can be correlated and can interact to define the niche.

## Methods

### General approach

We assume a variable is “important” if a model has high predictive accuracy and if the predictions are highly sensitive to changes in values of that variable (Meinshausen & Bühlmann 2010). To measure sensitivity of the model to changes in the variable, we use the permute-after-calibration test (Breiman 2001) which compares unpermuted and permuted predictions. We compared unpermuted and permuted predictions by calculating for each the Continuous Boyce Index (CBI), which indicates how well a model’s predictions serve as an index of the probability of presence (Boyce et al. 2002; Hirzel et al. 2006), and the area under the receiver-operator curve (AUC), which indicates a model’s ability to differentiate between presence and non-presence sites. We also calculated the correlation between unpermuted and permuted predictions (COR; Breiman 2001), which reflects differences between the two sets of predictions. Increasing the effect of a variable in a model should decrease CBI, AUC, or COR relative to an unimportant variable.

To simulate species’ distributions, we defined a generative function of one or two variables on the landscape, then used it to produce a raster of the probability of occurrence. For each cell, true occupancy was determined using a Bernoulli draw with the probability of success equal to the simulated probability of presence (Meynard & Kaplan 2013). We then calibrated and evaluated three SDM algorithms: generalized additive models (GAMs; Wood 2006), Maxent (Phillips et al. 2006), and boosted regression trees (BRTs; Elith et al. 2008). Full details of model calibration are presented in Appendix 1. Briefly, (unless otherwise stated) SDMs were calibrated using 200 occurrences and 10000 (Maxent and GAMs) or 200 (BRTs) background sites (Baret-Massin et al. 2012). Predictive accuracy and inferential capacity were evaluated using 200 distinct test occurrences plus either 10,000 background sites (CBI, AUC_bg_, and COR_bg_) or 200 absences (AUC_pa_ and COR_pa_; Meynard et al. 2019). In addition to the permute-after-calibration test, for each of the experiments we also evaluated algorithm-specific measures of variable importance: AIC-based variable weighting for GAMs (Burnham & Anderson 2002); contribution, permutation, and change-in-gain tests for Maxent (Phillips & Dudík 2008); and deviance reduction for BRTs (Elith et al. 2008). To streamline discussion, we present results only for Maxent and CBI in the main text (Appendices 3-9 present the complete set of results).

In our simulations there is no process-based spatial autocorrelation in species occurrences arising from dispersal, disturbance, or similar processes. As a result, occurrences at sites are statistically independent of one another regardless of their proximity, which obviates the need to use geographically distinct training and test sites. This is a convenience that is unlikely to be met in real-world situations where robust inference requires test data that is geographically and/or temporally as independent as possible from training data (Roberts et al. 2017; Fourcade et al. 2018).

For each level of a factor we manipulated in an experiment (e.g., landscape size), we generated 100 landscapes with training and test data sets, then calibrated and evaluated GAM, Maxent, or BRT models on each set. As a benchmark, we used an “omniscient” (OMNI) model which had the same variables and parameter values as the generative model used to create the species’ probability of occurrence (Meynard et al. 2019). We evaluated the OMNI model with the same set of test data used to evaluate the SDMs. Since OMNI does not use training data variability in its results is due solely to stochastic differences in test sites between iterations. To generate predictions from permuted variables, for a given level of a factor, data iteration, and SDM algorithm, we created 30 permutations of each variable, calculated test statistics (CBI, AUC, and COR) for each, then took the average test statistic value across these 30 sets.

We know a priori that levels of a factor are different, so our interest is in the effect size of each factor level (White et al. 2014), which can be discerned by eye. For a given test statistic (CBI, AUC, or COR), we assessed the capacity of the permute-after-calibration test to assess *discrimination* (qualitative differences between variables with different influence) and *calibration* (how well the distribution of the SDM’s test statistic matches that of the test statistic generated using the unbiased OMNI model). We designate a test as having “reliable discrimination” if there is complete lack of overlap between the inner 95% of the distribution of the test statistic between the permuted and unpermuted predictions across the 100 iterations. We designate a test as having “reliable calibration” by comparing the distribution of the test statistic between the SDM and OMNI: the range of the SDM’s inner 95% of values across data iterations are within ±10% of OMNI’s range, and the SDM’s median value is within the 40th and 60th percentile of OMNI’s median value. Under this definition, neither CBI, COR_pa_, nor COR_bg_ were ever well-calibrated, although in a few cases AUC_pa_ and AUC_bg_ yielded well-calibrated outcomes (Appendices 5 and 6). Hence, hereafter we focus on discrimination accuracy.

### Experiment 1: Simple scenario

In the simplest scenario we assumed the species’ probability of occurrence is determined by a logistic generative function of a single TRUE variable:

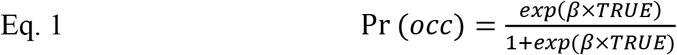

where *β* represents the strength of the response. We set *β* = 2 which produced in a moderate gradient in the probability of presence across the landscape. The TRUE variable has a linear gradient in geographic space ranging from −1 to 1 across a square landscape 1024 cells on a side (Appendix 2 Fig. S1). As a result, the species’ probability of occurrence is symmetrically distributed with an inflection point midway across the landscape. For each of the 100 data iterations, a Maxent model was trained using values of TRUE plus values of a spatially random FALSE variable with the range (−1, 1). The FALSE variable represents a variable “mistakenly” assumed to influence the species’ distribution.

### Experiment 2: Training sample size

Next, we examined how the number of occurrences used in the training sample affects estimates of variable importance. We used the same landscape and probability of occurrence as in *Experiment 1*. The number of training occurrences was varied across the doubling series 8, 16, 32, …, 1024, but the number of test sites was kept the same (200 occurrences and 10,000 background sites).

### Experiment 3: Prevalence

We then explored the effects of prevalence (mean probability of occurrence). The species responded to TRUE as per a logistic function,

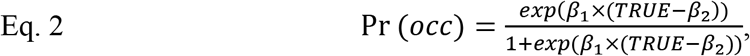

where non-zero values of the offset parameter *β*2 shift the range across the landscape, thereby altering prevalence (Appendix 2 Fig. S2). This allows us to manipulate prevalence while not changing study region extent, which is almost impossible in real-world situations. We set *β*1 equal to 2 and chose values of *β*2 that varied prevalence in 9 steps from 0.05 to 0.95.

### Experiment 4: Study region extent

In real-world situations enlarging a study region typically decreases prevalence (Anderson & Raza 2010) while also increasing the range of variability in environmental variables (VanDerWal et al. 2009). In this experiment we isolate the effect of increasing the extent of the study region on the range of environmental variation in the TRUE variable. Landscape size was varied along the doubling series 128, 256, 512, …, and 8192 cells on a side, each matched with an increasing range of the TRUE variable from (−0.125, 0.125) for the smallest landscape to (−8, 8) for the largest (Appendix 2 Fig. S3). FALSE was spatially randomly distributed and had the range (−1, 1). The species responded to TRUE as per a logistic function (Eq. 1). As a result, increasing extent did not affect prevalence.

### Experiment 5: Spatial resolution and autocorrelation of environmental data

In this experiment we explored the effects of spatial autocorrelation and spatial resolution (grain size) of environmental data on estimates of variable importance. We began with a linear TRUE and random FALSE variable distributed across a landscape with 1024 cells on a side. When unperturbed, the linear gradient of the TRUE variable has a high degree of spatial autocorrelation because cells close to one another have similar values. To manipulate spatial autocorrelation, we randomly swapped values of a set proportion of cell values (no cells, or one third, two thirds, or all of the cells). Swapping values reduces spatial autocorrelation in the TRUE variable because cells with dissimilar values were more likely to be close to one another (Appendix 2 Fig. S4). FALSE had a level of spatial autocorrelation no different from random, regardless of swapping. We assumed the species responded to the environment at the “native” 1/1024th scale of the landscape. Probability of occupancy was modeled with Eq. 1, and training and test sites were sampled at this “native” resolution. In some of the simulations we changed the spatial resolution of the environmental data presented to the SDMs by resampling the environmental rasters to a finer resolution with 16,384 cells or to a coarser resolution with 64 cells on a side using bilinear interpolation (sample size was kept the same regardless of resolution). In summary, we created a landscape, (possibly) swapped cells, modeled the species’ distribution and located training and test sites at the “native” scale, then assigned environmental values to sites based on the landscape at the (possibly) resampled resolution. This recreates a realistic situation where occurrences represent the scale of the true response but environmental data used to predict the response are available at a (potentially) different resolution. We explored all combinations of grain size (cell size of 1/16,384th, 1/1024th, and 1/64th of the landscape’s linear dimension) and spatial autocorrelation (swapping no cells, or one third, two thirds, or all of the cells; Appendix 2 Fig. S4).

### Experiment 6: Collinearity between environmental variables

Next, we explored the effects collinearity (correlation) between FALSE and TRUE predictors. As before, the species had a logistic response (Eq. 1) to TRUE, which has a linear gradient across the landscape. In contrast to previous experiments, FALSE also has a linear trend which is rotated relative to the gradient in TRUE to alter the correlation between the variables. We used a circular landscape to ensure no change in the univariate frequencies of the variables with rotation. The two variables are uncorrelated when their relative rotation is 90°, and positive or negative if less than or more than 90°, respectively (Appendix 2 Fig. S5). We used rotations of FALSE relative to TRUE from 22.5 to 157.5° in steps of 22.5°, which produced correlations between the two variables ranging from −0.91 to 0.91. Both variables had values in the range (−1, 1).

### Experiments 7, 8, and 9: Two influential variables

In the final experiments the species’ niche was shaped by two influential variables, T1 and T2, which both have linear gradients on a circular landscape and values in the range (−1, 1). The species responds to T1 and T2 as per a Gaussian bivariate function:

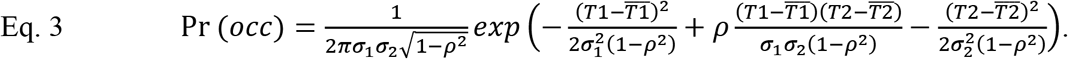

Niche breadth in T1 and T2 is determined by *σ*_1_ and *σ*_2_. Importantly, decreasing *σi* increases the degree to which a variable restricts distribution, meaning that *σi* and variable importance are inversely related. Changing *σ*_1_ and *σ*_2_ also alters the distribution of the species (Appendix 2 Fig. S6). Parameter *ρ* determines the degree of “niche covariance,” or interaction between variables in shaping the niche.

In *Experiment 7* we examined the effects of niche breadth by varying *σ*_1_ and *σ*_2_ across all combinations of 0.1, 0.3, and 0.5. We set niche covariance *ρ* = 0 and kept T1 and T2 uncorrelated on the landscape. In *Experiment 8* we investigated the effects of niche covariance by varying *ρ* from −0.75 to 0 to 0.75. We used intermediate niche breadth (*σ*_1_ and *σ*_2_ = 0.3) and kept T1 and T2 uncorrelated. Finally, in *Experiment 9* we explored all combinations of niche breadth (varying *σ*_1_ and *σ*_2_ from 0.1, 0.3, to 0.5), niche covariance (varying *ρ* from −0.5, 0, to 0.5), and collinearity between T1 and T2 on the landscape (varying *r* from −0.71, 0, to 0.71).

### Reproducibility

We created the R (R Core Team 2018) package [URL redacted for double-blind peer review] for generating virtual species and assessing variable importance (available at [URL redacted]). Code for the experiments and figures in this article is available at [URL redacted]. The package and code for the experiments depend primarily on the *dismo* (Hijmans et al. 2017), *raster* (Hijmans 2018), and [redacted] packages for R.

## Results

In each experiment we assessed six metrics (Box 1). Results from OMNI serve as a benchmark for the SDM because they represent the best an SDM could be expected to do given only variation in test data. Thus, when unpermuted predictions from OMNI have poor predictive accuracy, or when OMNI with permuted predictions fails to discriminate between variables, the SDM should also fail. Importantly, results for a given level of a manipulation represent outcomes across multiple data iterations, each of which typically spanned a much smaller range (during permutation) than the full set of models. Modelers typically have just one set of data for a species, so variation for a single data instance will underestimate the uncertainty inherent in the data sampling process.

### Experiment 1: Simple scenario

OMNI with unpermuted variables had high predictive accuracy (Fig. 1a). OMNI TRUE and FALSE permuted did not overlap, meaning that the variables could be successfully differentiated. Maxent performed similarly, although with more variation around TRUE permuted compared to OMNI.

**Figure 1.**
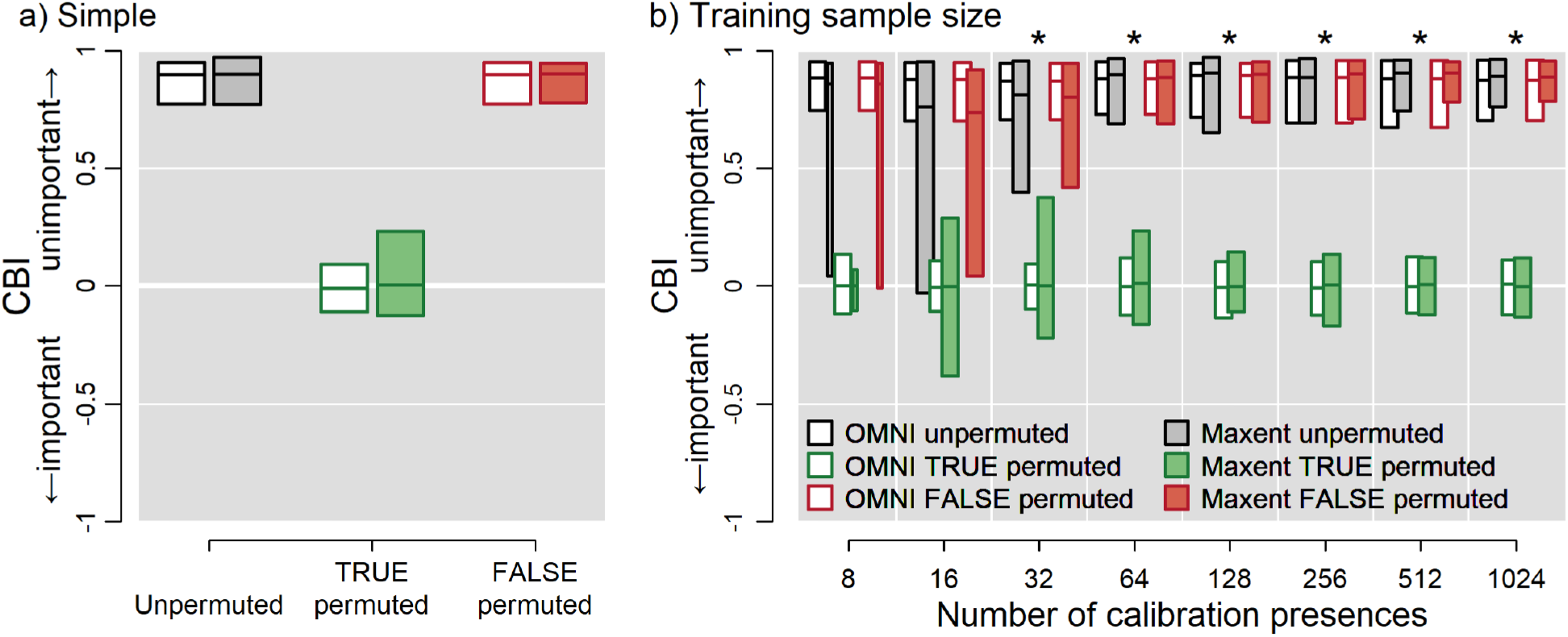
(a) *Experiment 1: Simple scenario*. The species’ range is influenced solely by a TRUE variable, but data on an uninfluential FALSE variable is also “mistakenly” presented to models. Maxent successfully discriminates between the two variables (no overlap between green and red bars). See Box 1 for further interpretation. (b) *Experiment 2: Sample size*. Maxent cannot differentiate between TRUE and FALSE at *n* <32. Asterisks indicate cases where OMNI and Maxent reliably discriminate between TRUE and FALSE. Bar width is proportional to the number of Maxent models that are more than intercept-only models (typically *n* = 100; CBI cannot be calculated if there is no variation the response). See Box 1 for further guidance on interpretation.

### Experiment 2: Training sample size

OMNI does not use training data, so always correctly discriminated TRUE from FALSE regardless of training sample size (Fig. 1b). Maxent unpermuted performed as well as OMNI unpermuted when sample size was ≥64, but below this Maxent had much greater variability than OMNI and was unable to reliably discriminate between TRUE and FALSE at *n* <32. At the smallest sample size (*n* = 8), Maxent often yielded intercept-only models that could not be used to calculate CBI, which requires variation in predictions for calculation.

### Experiment 3: Prevalence

Increasing prevalence (mean probability of occurrence) reduced unpermuted OMNI’s predictive accuracy and increased variability. As a result, OMNI often performed no better than random when prevalence was ≥0.85 (Fig. 2a) and could not discriminate between TRUE and FALSE. Maxent was qualitatively the same, although variation in permuted and unpermuted CBI was greater at high prevalence than in OMNI (Fig. 2a).

**Figure 2.**
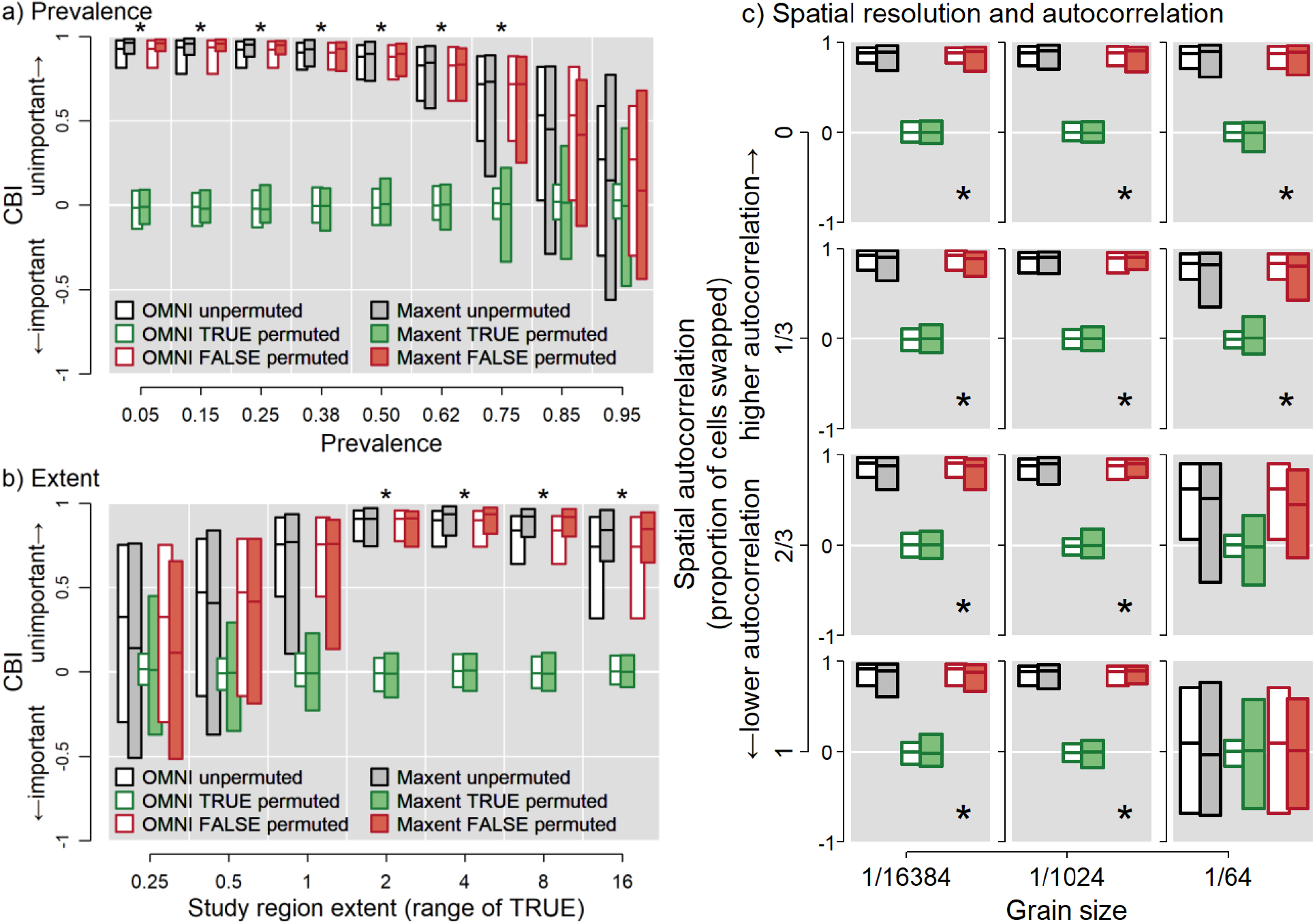
(a) *Experiment 3: Prevalence*—Effect of prevalence on inferential power. Neither Maxent nor OMNI reliably discriminate when prevalence is >0.75. (b) *Experiment 4: Study region extent*—Maxent measures variable importance most reliably when the study region is large enough to encompass sufficient environmental variation. Study region extent is indicated by the range of the TRUE variable which is proportional to the size of the landscape. (c) *Experiment 5: Spatial resolution and autocorrelation*—The species perceives the environment at a “native” resolution such that cells are 1/1024th of the linear dimension of the landscape. Environmental data were downscaled to cells 1/16,384th on a side or upscaled to 1/64th on a side. Spatial autocorrelation was decreased by randomly swapping cell values of 1/3, 2/3 or all of the cells. Maxent fails when resolution is coarse and autocorrelation low. In all panels asterisks indicate both OMNI and Maxent reliably discriminate between TRUE and FALSE. See Box 1 for further guidance on interpretation.

### Experiment 4: Study region extent

Increasing landscape extent (the range of the TRUE variable) caused predictive accuracy of OMNI unpermuted to peak at intermediate extents where the range of TRUE was from 2 to 4, which occurs when the landscape was from 1024 to 2048 cells on a side (Fig. 2b). OMNI failed to reliably discriminate between TRUE and FALSE on the smallest landscapes (range of TRUE ≤0.5). Maxent was less reliable, failing when the range of TRUE was ≤1 (Fig. 2b). The worsening performance of OMNI unpermuted at large extents might be due to the sensitivity of CBI to test presences that are located in areas with a very low probability of presence and to potential underfitting of Maxent (Appendix 10).

### Experiment 5: Spatial resolution and autocorrelation of environmental data

The “true” importance of TRUE and FALSE is indicated by OMNI at the “native” resolution of 1/1024 (middle column of subpanels in Fig. 2c). Results for OMNI at the other resolutions represent the outcome that would be obtained if a modeler had perfect knowledge of the species’ response to the environment but only had data available at finer or coarser resolutions. Both OMNI and Maxent reliably discriminated between TRUE and FALSE at the “native” 1/1024th resolution and at the finer 1/6384th resolution, plus at the coarser 1/64th resolution when spatial autocorrelation was high. However, when the environmental data was coarse (1/64th scale) and spatial autocorrelation low (proportion swapped two-thirds or 1), both OMNI and Maxent unpermuted overlapped or nearly overlapped 0, indicating some models performed no better than random. Neither model could reliably discriminate between TRUE and FALSE in these cases (cf. Meynard et al. 2019).

### Experiment 6: Collinearity between environmental variables

OMNI completely ignores the FALSE variable and thus had high performance regardless of the magnitude of correlation between TRUE and FALSE (Fig. 3). Although Maxent unpermuted was slightly more variable than OMNI, Maxent had fairly high predictive accuracy across the entire range of correlation between TRUE and FALSE. Maxent also reliably discriminated between TRUE and FALSE except when collinearity was high (|*r*| >0.71), although even small amounts of correlation caused TRUE permuted to vary more than OMNI TRUE permuted. The increasing range of Maxent FALSE permuted at high magnitudes of correlation indicate that Maxent sometimes used information in the FALSE variable (Fig. 3).

**Figure 3.**
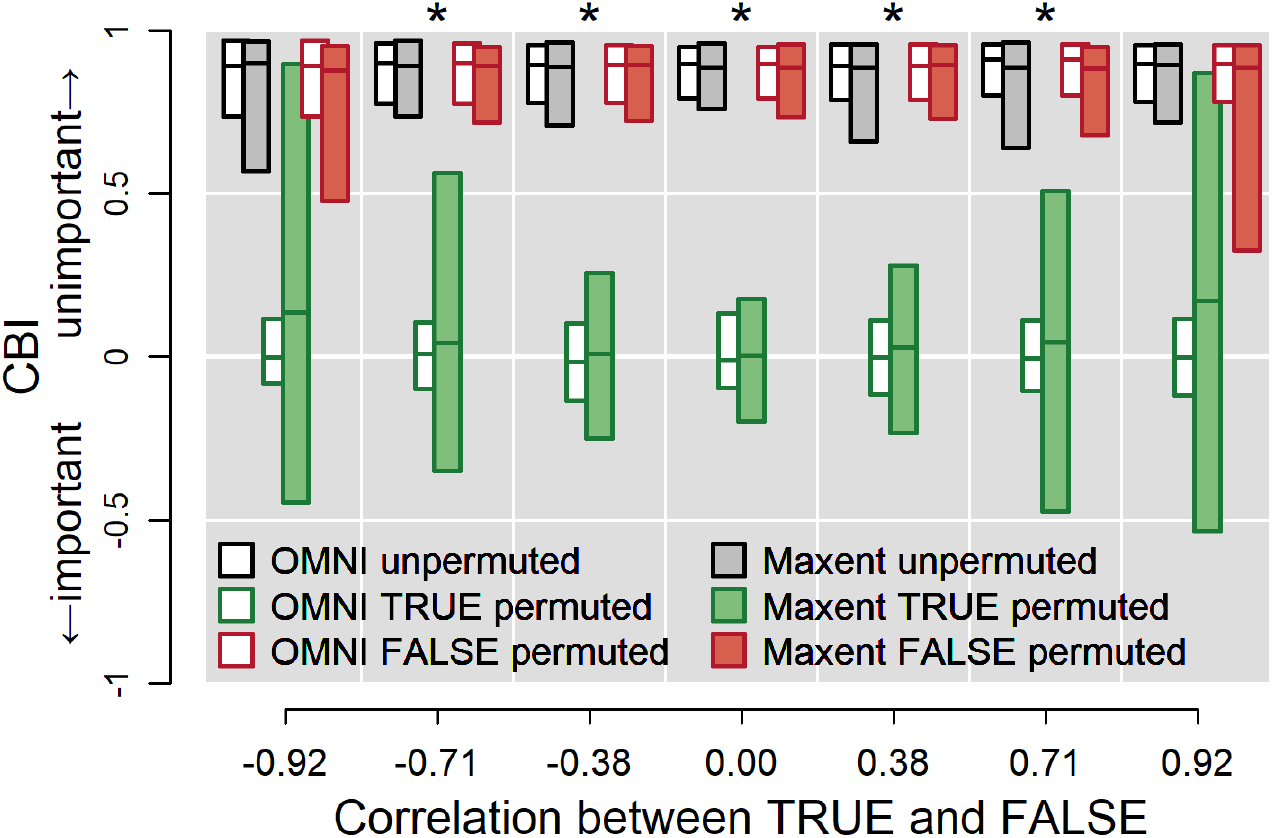
*Experiment 6: Collinearity*. The species’ range is determined by a single TRUE variable, but Maxent is presented data on this variable plus a correlated FALSE variable. FALSE is increasingly used by Maxent as the magnitude of correlation increases. Asterisks indicate both OMNI and Maxent reliably discriminate between TRUE and FALSE. See Box 1 for further guidance on interpretation.

### Experiments 7 through 9: Two influential variables

In *Experiment 7* we manipulated niche breadth of two influential variables while keeping all other factors “off” (no niche covariance, no correlation). In cases where both variables had equal influence on the niche (*σ*_1_ = *σ*_2_), OMNI unpermuted had high predictive accuracy across all combinations of niche breadth in T1 and T2 (median CBI ranging from 0.89 to 0.93; Fig. 4a). Maxent unpermuted also had high predictive accuracy. Permuting a variable for which niche breadth is narrow should reduce CBI more than when it is broad, which is what we observed with OMNI. For example, changing niche breadth from broad (*σ*_1_ = *σ*_2_ = 0.5) to medium (*σ*_1_ = *σ*_2_ = 0.3) to narrow (*σ*_1_ = *σ*_2_ = 0.1) reduced median CBI from 0.68 to 0.63 to 0.46, respectively (similar values were achieved for T2). However, Maxent did not show a monotonic decline; respective values were 0.68, 0.76, and 0.64 (Fig. 4a). (Permuting T2 yielded similar anomalies). Thus, Maxent always underestimated the importance of T1 and T2 when niche breadth was moderate or narrow and variables acted equally.

**Figure 4.**
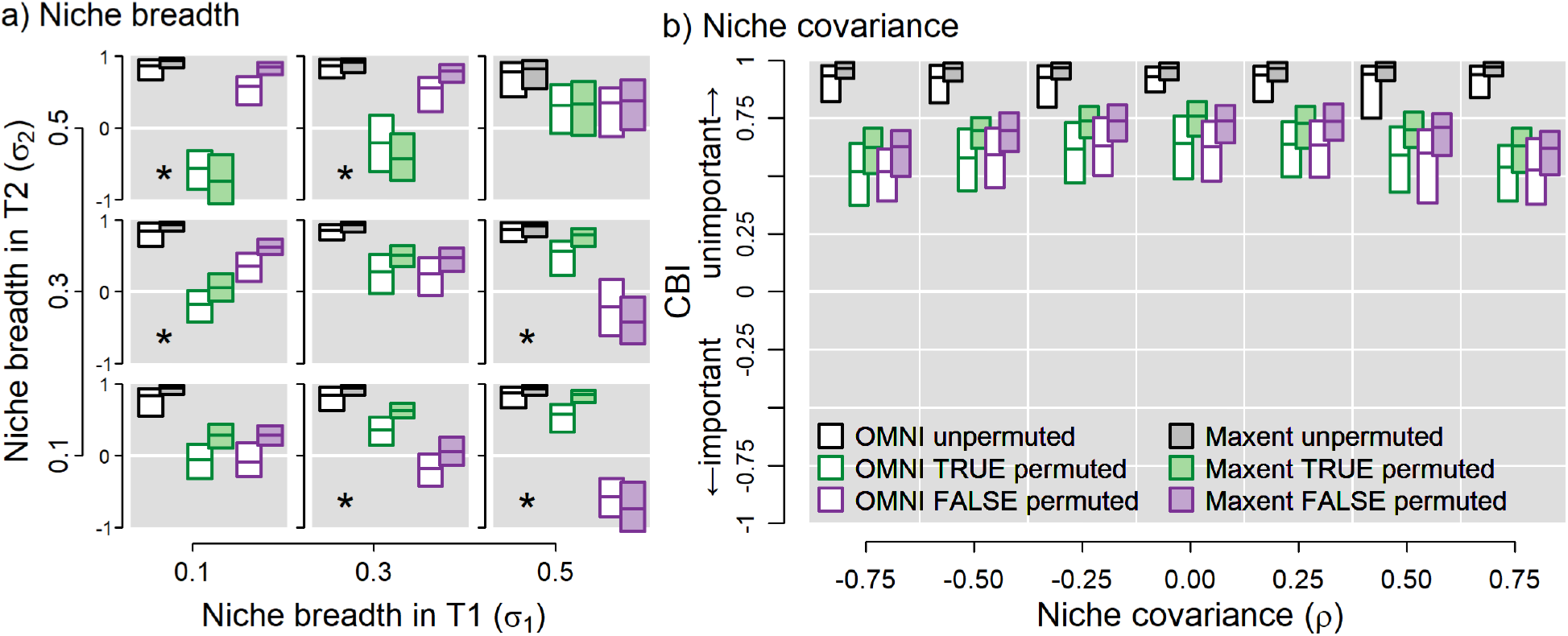
(a) *Experiment 7: Niche breadth*. Each subpanel represents results from modeling a species on a landscape with two influential variables, T1 and T2, with niche breadth set by *σ*_1_ and *σ*_2_. Narrower niche breadth increases limitation by that variable so should yield lower CBI when the variable is permuted. The y-axis on each subpanel represents CBI. The case shown here is for a landscape with no correlation between variables (*r* = 0) and no niche covariance (*ρ* = 0). Asterisks indicate both OMNI and Maxent reliably discriminate between T1 and T2. (b) *Experiment 8*: *Niche covariance*. Variables interact to define the niche. The case shown here is for a landscape with no correlation (*r* = 0) with moderate and equal niche breadth in both variables (*σ*_1_ = *σ*_2_ = 0.3). See Box 1 for further interpretation.

In cases where variables had asymmetrical influence (*σ*_1_ ≠ σ_2_), OMNI unpermuted had high predictive accuracy, and permuting T1 or T2 caused CBI to decrease monotonically with decreasing niche breadth in the respective variable (Fig. 4a). Maxent always reliably discriminated between T1 and T2 (i.e., no overlap between distributions of permuted CBI for T1 and T2). However, Maxent tended to overestimate the importance of the more influential variable. Estimates of importance for the less influential variable tended to be more uncertain compared to the more influential variable.

In *Experiment 8* we altered niche covariance (interaction between variables) but kept niche breadth and correlation between variables fixed. Increasing the magnitude of covariance increased the actual and estimated importance of the variables even though niche breadth was held constant (*σ*_1_ = *σ*_2_ = 0.3; Fig. 4b). On average, Maxent underestimated the importance of the variables at all levels of covariance.

In *Experiment 9* we examined all possible combinations of niche breadth and covariance and correlation between variables. Results were qualitatively similar to the preceding two experiments (Appendix 4 Fig. S9).

### Results using AUC and COR, algorithm-specific tests, and different algorithms

Results using the permute-after-calibration test paired with AUC_pa_, AUC_bg_, COR_pa_, and COR_bg_ and algorithm-specific tests are summarized in Appendix 3 and presented in detail in Appendices 4 through 9, so are only recapitulated here. We found notable interactions between model algorithm and the metric used by the permute-after-calibration test. For example, GAMs were capable of discriminating between TRUE and FALSE at training sample sizes as low as 8 (though few models converged) when using CBI or COR (either variant), but required 128 or more occurrences when using AUC (either variant). However, results using AUC were well-calibrated at large sample sizes. Across all experiments, there was no best test statistic, although AUC was much less variable than CBI, and COR was much more variable. Maximum values of AUC for unpermuted OMNI models were always substantially <1.

Although Maxent and GAMs performed differently in most experiments, neither consistently outperformed the other across experiments. GAMs tended to show less variation in simple cases (TRUE versus FALSE) but more in complex cases (TRUE versus TRUE). In all experiments BRTs had much greater variation and thus less reliable discrimination than the other two algorithms.

In general, algorithm-specific tests were less robust to challenging situations than the permute-after calibration-test. For example, Maxent’s change-in-gain test was only able to discriminate between TRUE and FALSE when sample size was ≥128 (Appendix 9 Fig. S2), but minimum necessary sample size was 64 using COR or Maxent’s permutation or contribution tests, and just 32 when using CBI or AUC.

## Discussion

Our objective was to delineate the minimal necessary conditions under which species distribution models and ecological niche models can be used to infer variable importance. We found that the permute-after-calibration test was capable of discriminating between variables under many situations, but results typically differed from expectations established by an “omniscient” model. Notably, high predictive accuracy did not necessarily connote high inferential capacity. However, situations that are challenging for generating SDMs with high predictive accuracy were also challenging for estimating variable importance (results summarized in Table 1): small sample size, high prevalence, low spatial extent (low environmental variability), high collinearity, and using environmental data that is coarser than the perceptual scale of the species when spatial autocorrelation is low. When more than one variable shaped a species’ distribution, SDMs were able to discriminate between two variables with different influence, but mis-calibrated importance when variables acted equally. Interactions between variables in how they shaped the niche had little effect on discriminatory power. In general, factors that shape inference can be classified into those that are extrinsic to the species (e.g., choice of modeling algorithm, sample size) and those that are intrinsic to the species (e.g., niche breadth). Extrinsic factors are at least nominally under the control of the modeler and thus offer the potential for amelioration, whereas confounding intrinsic factors likely require development of new techniques and robust data to control for their influence. We structure the discussion around what modelers typically can control and what they typically cannot.

**Table 1.**
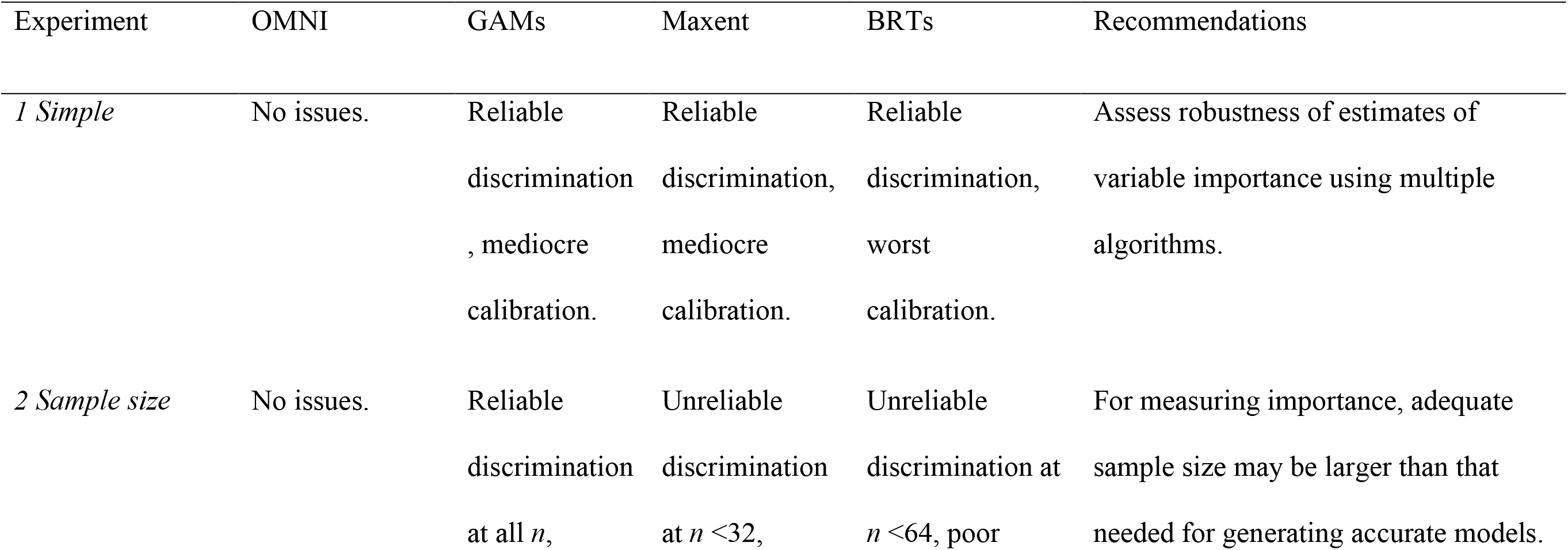
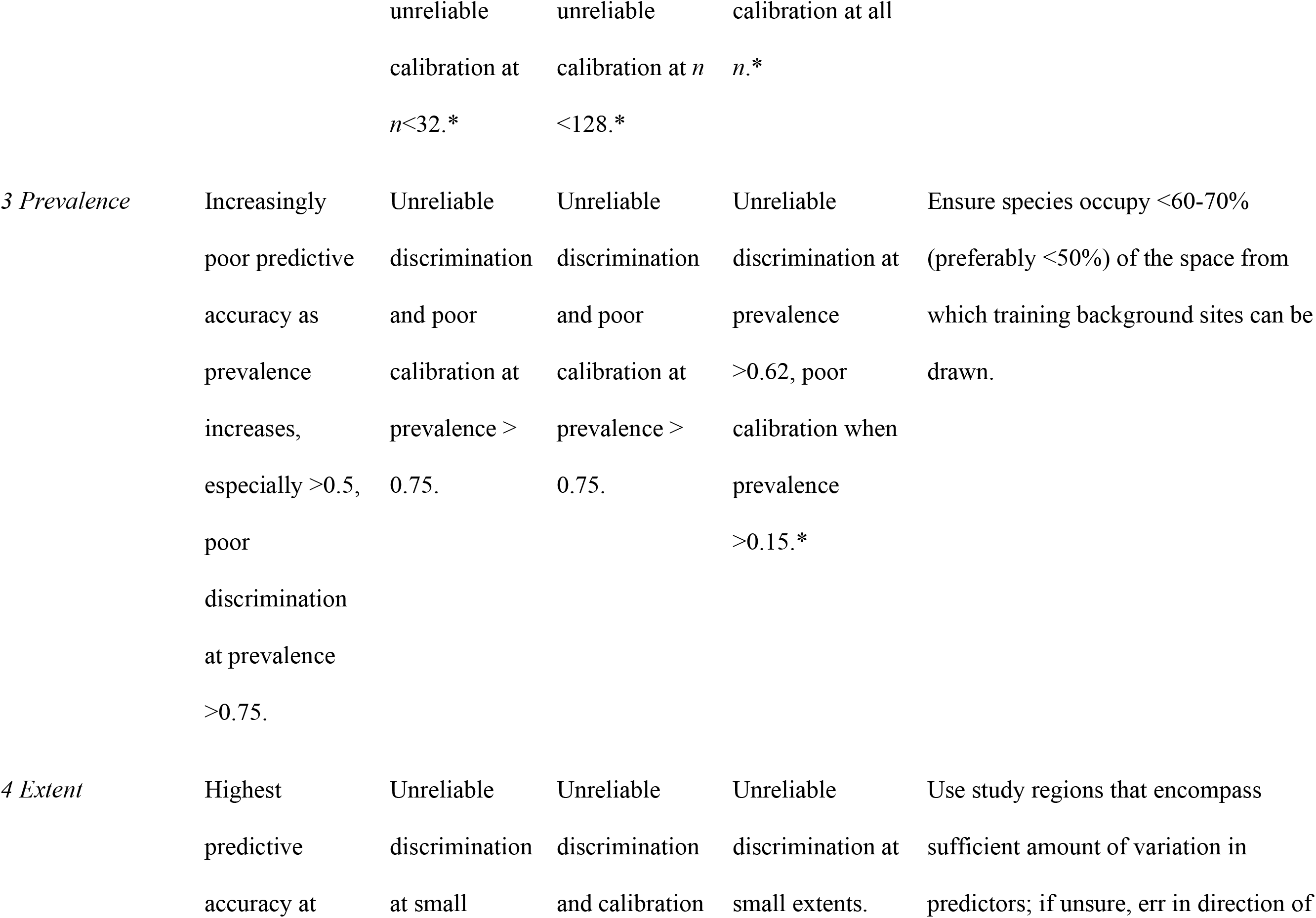
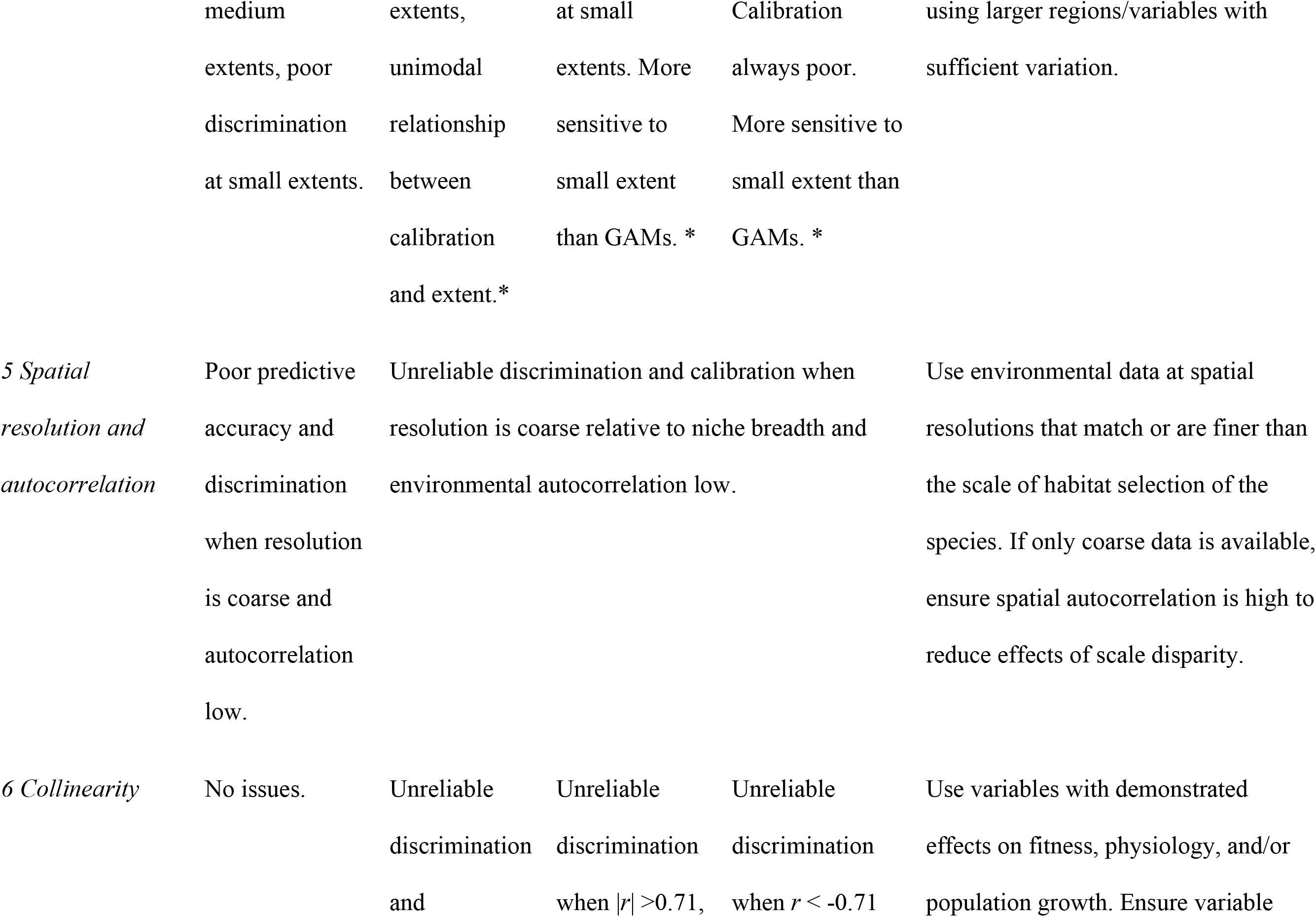
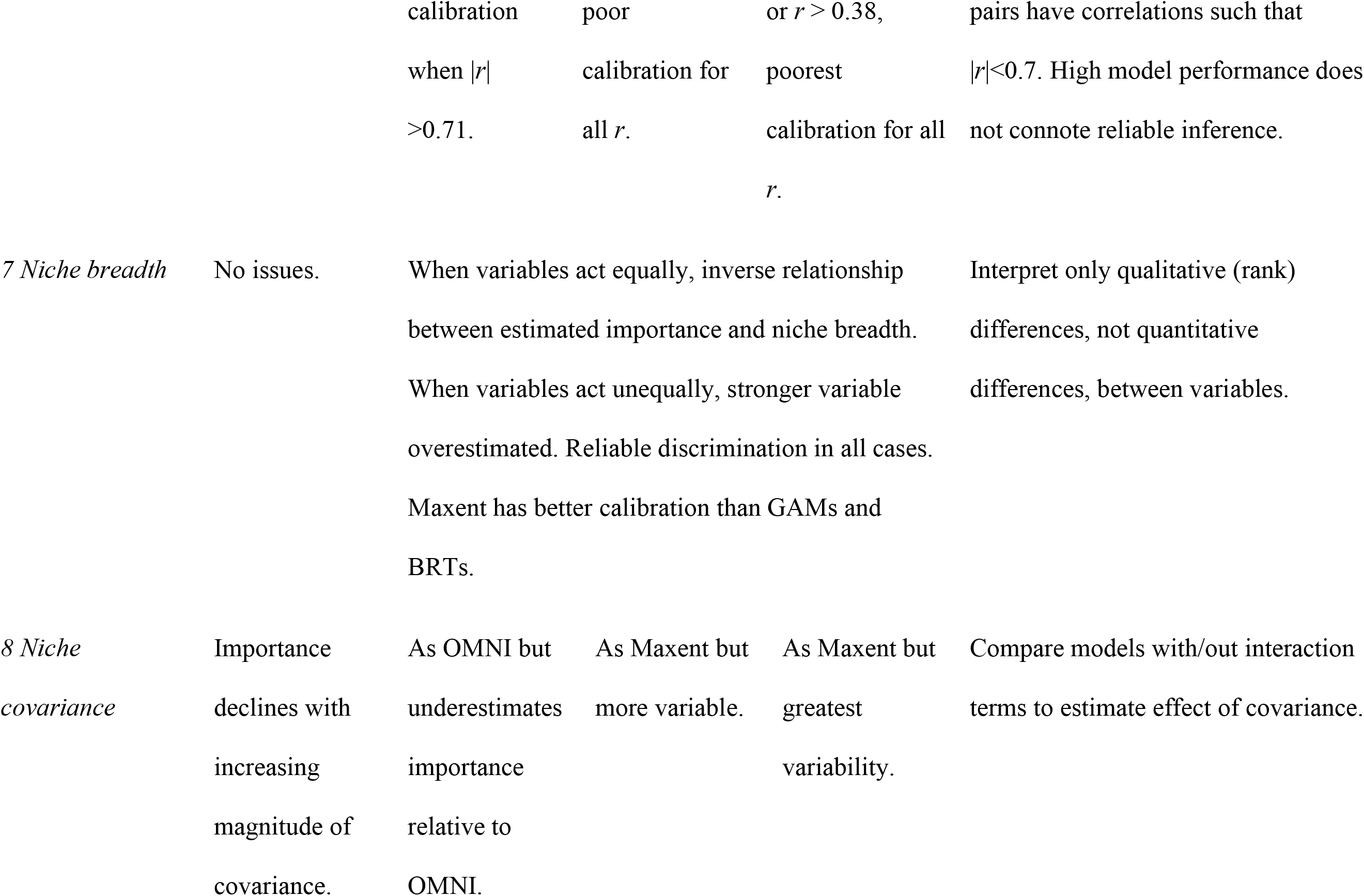
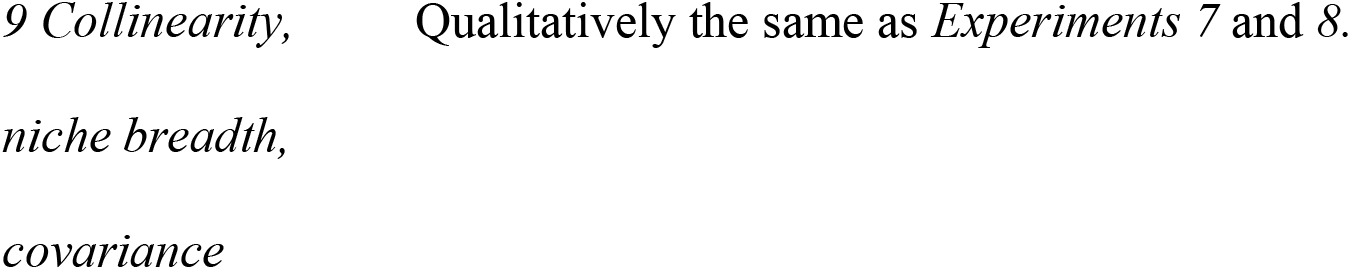
Summary of results across the nine simulation experiments for the permute-after-calibration test. Discrimination refers to the ability to differentiate between two variables with different influence. Few tests met our quantitative standard for reliable calibration, so here “good calibration” refers to a subjective comparison between the tests result and the results using an “omniscient” model (i.e., OMNI). Asterisks indicate the algorithm sometimes had convergence issues or yielded intercept-only models which disallowed calculation of test statistics. Results in this table are particular to using the permute-after-calibration test paired with CBI. Thresholds using other metrics or algorithm-specific tests were sometimes different, but recommendations remain unchanged.

### Model algorithm and inferential method

The reliability of tests of variable importance depended on the algorithm, type of test, and metric used to evaluate the test. We found GAMs and Maxent had much less variability and thus greater discriminatory capacity than BRTs, despite extensive efforts to tune BRTs (Appendix 1). Choice of modeling algorithm is one of the largest contributors to variation in predictive capacity (Dormann et al. 2008; Barbet-Massin et al. 2012; Rapacciuolo et al. 2012), with no one algorithm necessarily best for all species (Qiao et al. 2015). Our results show that model choice also affects inferential power and thus underscores the importance of evaluating variable importance using multiple algorithms.

We also found inferential capacity varied by the nature of the test (permute-after-calibration test versus algorithm-specific tests) and associated test metric (e.g., CBI, AUC, change in Maxent’s gain). Depending on the situation and algorithm, the choice of test metric (CBI, AUC, COR) affected the reliability of the permute-after-calibration test, but no one metric consistently out-performed the others in discrimination capacity. AUC was occasionally better-calibrated than CBI and COR, especially when paired with GAMs, but also had less reliable discrimination in these same circumstances. In contrast, algorithm-specific tests were less robust to challenging circumstances than the permute-after-calibration test (Appendix 3).

Despite the better performance of the permute-after-calibration test, care needs to be used to ensure the metric with which it is paired provides unconfounded interpretation. Namely, tests of variable importance can only reliably indicate differences between variables if the original model (i.e., with unpermuted predictions) has high predictive accuracy (Meinshausen & Bühlmann 2010). Neither AUC nor COR are capable metrics in this respect. In particular, maximum AUC (either variant) is typically depressed well below 1 (Jiménez-Valverde et al. 2013; Smith 2013a), which is evident even in our simplest scenario (*Experiment 1*) where median unpermuted AUC_bg_ and AUC_pa_ for the OMNI model was only ~0.78 and ~0.64, respectively. Likewise, COR does not indicate the predictive capacity of a model. Given these considerations, we recommend 1) employing multiple modeling algorithms that can be compared using 2) algorithm-independent tests like the permute-after-calibration test; and 3) employing metrics that can be objectively interpreted as measures of predictive accuracy and that are not known to be influenced by study-specific aspects like prevalence or sample size (Jiménez-Valverde et al. 2013; Jiménez-Valverde 2020; Smith 2013a).

### Sample size

We found that inferential power was compromised when training sample size was between 8-128, depending on the algorithm and test. However, owing to the fact that simulations ensured other conditions were optimal (e.g., no dispersal, perfect detection), we expect the actual minimum sample size for reliable application real-world situations to be larger, especially given biases in data collection (Feeley & Silman 2011). Although new techniques amenable to modeling rare species (Lomba et al. 2010; Breiner et al. 2015) might be able to lower the threshold sample size necessary for predictive accuracy, small samples can still induce spurious correlations between predictors (Ashcroft et al. 2011) and may not adequately capture the full extent of species’ environmental tolerances (Feeley & Silman 2011). On these bases, we expect that minimal sample size for reliable inference will be several times larger than sample size necessary for generating models with high predictive accuracy.

### Spatial scale: Extent, prevalence, resolution, and autocorrelation

We found inferential power declined rapidly when prevalence was >0.5 (Fig. 2a) and when the study region extent was too small to encompass sufficient environmental variation to distinguish occurrences from non-occurrences (Fig. 2b). In real-world situations, decisions regarding spatial extent of the study region typically affect prevalence, the range of environmental variability in training data, and degree of spatial autocorrelation and collinearity between predictors (Seo et al. 2008; VanDerWal et al. 2009; Lauzeral et al. 2013). Thus, there are likely interactions and cascading effects of decisions about scale that are not apparent in our results. For example, study extent can interact with spatial autocorrelation to affect the variables that appear important in a model (VanDerWal et al. 2009; Connor et al. 2017).

We also found that inference was compromised when spatial resolution was coarser than the species’ scale of perception and spatial autocorrelation was low (Fig. 2c). Best practices recommend using environmental data at a resolution matching the scale of species’ response to the environment (Mertes & Jetz 2018; Araújo et al. 2019), although scale mismatch can be ameliorated when spatial autocorrelation is high (Fig. 2c; Moudrý & Šímová 2012; Mertes & Jetz 2018). To date, the finest resolution climate data with global-scale coverage has a resolution on the order of ~1 km (Fick & Hijmans 2017; Karger et al. 2017), which is much larger than the scale of perception of the environment of most sessile and many mobile organisms. Hence, scale mismatch will likely remain a problem for many studies.

Based on our results, we recommend at the minimum ensuring the region from which background sites are drawn is large enough to encompass sufficient environmental variation and that the species occupies less than about half the landscape. Likewise, when fine-scale environmental data is not available, we recommend at least measuring spatial autocorrelation to assess the degree to which scale mismatch could confound inference (Naimi et al. 2014).

Modelers must be aware that the results of an inferential study will be dependent on all aspects of scale and that these aspects can interact to affect inference in ways not explored here (e.g., VanDerWal et al. 2009; Hanberry 2013; Connor et al. 2017). As a result, comparisons between inferential studies that vary in aspects of scale need to be made with these complications in mind.

### Collinearity

Our results indicate that inferential power is low when the magnitude of pairwise correlation is >0.7 (Fig. 3). Alarmingly, unpermuted predictions often had high predictive accuracy even when high collinearity caused them to mistakenly use information in the FALSE variable. This is surprising but supported by other work that finds using predictors with no actual relationship to a species’ occurrence can yield models as accurate when using “real” variables (Buklin et al. 2015; Fourcade et al. 2018). Thus, the predictive accuracy of a model is not a reliable indicator of its inferential capacity.

Of all of our findings, the inability of models to differentiate between influential and uninfluential correlated variables, yet produce seemingly accurate predictions is the most troubling (Warren et al. 2020). Environmental variables are often collinear (Jiménez-Valverde et al. 2009), so this is likely a very frequent challenge to successful inference. However, modelers have some means to modulate collinearity. The simplest solution is to simply select variables that have low pairwise-correlations (Dorman et al. 2013). Unfortunately, discarding correlated variables inherently assumes dropped variables have zero influence with absolute uncertainty. Another solution is to employ modeling algorithms with regularization or regularization-like-behavior, but the methods used here already do that (e.g., Maxent LASSO; Tibshirani 1996; Phillips et al. 2006) and were not entirely robust to collinearity (Dormann et al. 2013). A third potential solution may be to construct multiple models with different sets of relatively uncorrelated variables (Barbet-Massin et al. 2014; Petitpierre et al. 2017) then average variables’ importance across them.

### Qualities of the niche

Niche breadth and interactions between variables in shaping the niche are inherent to species and thus not under control by the modeler. Niche breadth has the most obvious relationship to variable importance since narrower environmental tolerance should translate into increased sensitivity of a model to changes in that variable. Surprisingly, we found that when two variables act to shape the niche equally, reducing niche breadth does not lead to a monotonic increase in estimated importance of the variables (Fig. 4a). Likewise, when two variables acted unequally to influence the niche, SDMs overestimated the importance of the more important variable. As a result, the relative difference between the permuted and unpermuted values of a test statistic should not be interpreted as a measure of the absolute importance of a variable. Rather, we recommend interpreting only qualitative (rank) importance (Barbet-Massin & Jetz 2014).

Niche covariance occurs when, for example, negative effects of high temperature on a species’ fitness can be offset by high values of precipitation (Smith 2013b). Interaction between niche dimensions rotates the orientation of the niche in niche space, thereby changing the range of environments occupied (Appendix 2 Fig. S7). As a result, the importance of niche covariance are not always obvious from examination of univariate niche breadth (Smith 2013b). We found that increasing the magnitude of niche covariance (*ρ* ≠ 0) increased actual and estimated importance compared to cases where variables acted independently (*ρ* = 0), but importance was still miscalibrated vis-à-vis an omniscient model (Fig. 4b). We did not find strong interactions between niche breadth, niche covariance, and collinearity for the range of each investigated here, although the simplicity of our simulations does not preclude different outcomes in real-world situations.

### Variable and model selection

Our work calls into question the common practice of using automated methods for variable and model selection (Barbet-Massin & Jetz 2014; Gobeyn et al. 2017; Guisande et al. 2017; Cobos et al. 2019). We found SDMs using uninfluential variables could still yield measures of predictive accuracy that qualified them as “good” models (Fig. 3; Buklin et al. 2015; Fourcade et al. 2018). As a result, we echo others’ recommendations to use expert-based selection of variables before conducting algorithmic-based screening (Mod et al. 2016; Gardener et al. 2019).

### Future directions

The simplified nature of our scenarios likely means that conditions we identify for reliable inference (Table 1) represent the minimum circumstances under which these tests perform robustly. Real-world applications will surely require larger sample sizes, less collinearity, smaller disparities in scale, et cetera, to be reliable. Given the many ecological questions informed by measures of variable importance, understanding the domain in which inferential tests can be trusted is a pressing priority. To this end, many questions must be addressed: How do tests of variable importance fare against real-world factors like biotic interactions, local adaptation, disturbance, dispersal, bias in sampling, realistic environmental variation, and so on? How does high-dimensional niche space affect inference? How do other tests of importance compare to the ones evaluated here? How does data type (presence/background versus presence/absence versus abundance) affect inference (Gábor et al. 2020)? Answering these questions will require expanding beyond the reductionist approach used in this work. One alternative is to simulate niches or distributions as realistically as possible, including realistically-structured landscapes, biotic interactions, dispersal limitation, and other ecological processes (e.g., Zurell et al. 2016; Warren et al. 2020), then apply a battery of procedures to identify situations that are conducive to measuring variable importance accurately (Groves & Lempert 2007 describe an analogous approach in policy analysis). Alternatively, the small subset of Earth’s species for which there is extensive field-based knowledge of range-shaping factors could be used to validate model-based inferences of variable importance (e.g., Angert et al. 2018).

### Conclusions

Our work represents the first systematic assessment of conditions under which SDMs can reliably estimate variable importance. The good news is that SDMs were able to discriminate between variables under conditions conducive to generating models with high predictive accuracy. The bad news is that high predictive accuracy did not necessarily connote reliable inference (cf. Warren et al. 2020). Factors extrinsic to species that can be influenced by modelers and factors intrinsic to species affect the ability to measure variable importance. Given the ubiquity with which these models are used to measure the importance of environmental factors in shaping species’ distributions and niches (Bradie & Leung 2017), we see a great opportunity and a great need for further research in this area.

## Acknowledgements

We wish to thank three anonymous reviewers and the subject editor who dedicated the unreimbursed time and attention to improve the manuscript. This work was supported by the [redacted] to [redacted].

## Online appendices

Appendix 1: Model calibration

Appendix 2: Maps of landscapes and species

Appendix 3: Summary of results for CBI, AUC, COR, and algorithm-specific tests

Appendix 4: Using AUC calculated with presences and absences (AUC_pa_) as a test statistic for the permute-after-calibration test of variable importance

Appendix 5: Using AUC calculated with presences and absences (AUC_bg_) as a test statistic for the permute-after-calibration test of variable importance

Appendix 6: Using the correlation test statistic calculated with presences and absences (COR_pa_) as a test statistic for the permute-after-calibration test of variable importance

Appendix 7: Using the correlation test statistic calculated with presences and absences (COR_bg_) as a test statistic for the permute-after-calibration test of variable importance

Appendix 8: Algorithm-specific metrics of variable importance

Appendix 9: Sensitivity of the Continuous Boyce Index to improbable test presences

## Notes

### Competing Interest Statement

The authors have declared no competing interest.

### Summary of Updates

Main text now only presents results for Maxent. Appendices now include all results for Maxent, GAMs, and BRTs. Added results using algorithm-specific tests of variable importance (in Appendices). Responded to reviewers' concerns. Significantly shortened manuscript.

