## Appendix 1 for "Testing the ability of species distribution models to infer variable importance"

### Appendix 1: Model calibration

[Author names withheld for double-blind peer review.]

For most experiments in the main text we evaluated how well three distribution modeling algorithms measured variable importance, but for a few experiments we added three more algorithms. This document describes the manner in which each algorithm was calibrated ("tuned") to best match the training data while not overfitting it. All models were calibrated using functions in the [redacted for double-blind peer review] package [redacted for double-blind peer review] which are called by routines in the [redacted for double-blind peer review] package [redacted for double-blind peer review] for the R Statistical Environment (R Core Team 2018) version 3.5.2. Unless otherwise stated, we used the default settings for model training functions. Both packages rely primarily on the *dismo* (Hijmans et al. 2018) and *raster* (Hijmans 2018) software packages. For a given iteration in a particular scenario all algorithms were calibrated with the same set of presence and background sites and evaluated against the same set of test sites.

Except for *Experiment 2* where we examine the effect of sample size, we randomly selected 200 presences for model calibration. Depending on the model algorithm, we also selected 10,000 (GAMs and Maxent) or 200 (BRTs) random background sites for model calibration (Barbet-Massin et al. 2012). For model evaluation, we selected 200 test presences that were distinct from the presences used to calibrate the models. These were paired with 10,000 background sites for calculation of CBI, CORbg, and AUCbg¬, or with 200 test absences for calculation of CORpa and AUCpa. For each level of a factor we manipulated in an experiment (e.g., differences in prevalence, magnitude of correlation between variables, etc.), we generated 100 iterations of presences, absences, and background sites. We then trained and evaluated 100 GAMs, Maxent, and BRT models, one per set. The permute-after-calibration test requires variable values to be permuted, so for a given level of a factor, iteration, and model algorithm, we created 30 permutations of each variable, calculated CBI, AUC, or COR, then took the average value of CBI, AUC, or COR across these 30 sets.

#### Generalized additive models (GAMs)

GAMs are a flexible adaptation of generalized linear models in which the predictors are first "smoothed" using splines or similar functions then used in their modified form as predictors (Wood 2006). The algorithm is able to use different smooths for different portions of a predictor's range, allowing a potentially highly flexible fit. GAMs were calibrated using the *trainGam* function in the [redacted for double-blind peer review] package. The function is designed to balance issues related to small sample sizes relative to the number of predictors available for modeling. The function first constructs single-term models using a spline smooth for each variable plus a single-term model that includes two-way interactions between each pair of variables using a Tensor-product smooth (Wood 2006). $AIC_c$ is calculated for each model (Burnham & Anderson 2002). The function then adds terms to a "full" model in order from lowest to highest $AIC_c$ such that there must be at least 10 presence data points per term (univariate or bivariate), up to a limit of 8 terms. Then, all possible models are constructed from the "full" model, and the one with the lowest $AIC_c$ selected as the "best" model with the condition that there must be at least 20 presences per term in this model. GAMs were trained with 10,000 randomly located background sites (Barbet-Massin et al. 2012). The total weight of presence and background sites was equalized to be the same (Maggini et al. 2006).

**Maxent**

The Maxent algorithm first estimates the probability of the environment across the landscape from a set of background sites then inverts this using Bayes theorem to estimate the probability of presence (or an index thereof) given the environmental conditions at a site (Phillips et al. 2006; Phillips & Dudík 2008). We used Maxent Version 3.3.3k (Phillips & Dudík 2008). During the time we implemented the scenario modeling a newer version of Maxent (version ≥3.4.x; Phillips et al. 2017) became available. However, we found that on occasion it produced errors rooted in source code of sub-dependent packages which we were unable to fix or have addressed. As a result, we chose to use the slightly older version, but with the default complementary log-log ("cloglog") output (function *predictMaxEnt* in the [redacted for double-blind peer review]package; [redacted for double-blind peer review]) used by the newer version of Maxent. The two model versions are exactly equivalent down to rounding error, except that the older version explores all possible breakpoints for threshold features, whereas the newer version uses a smaller set of breakpoints to speed computation and the newer version does not use threshold features by default (Phillips et al. 2017).

To find the optimal master tuning parameter which adjusts the "smoothness" of fit and the optimal set of functional "features" used by the model (linear, quadratic, threshold, hinge, and two-way interaction), we calculated models with all possible combinations of features (which always included the "linear" feature) and values for the master tuning parameter (from 0.5 to 3 in steps of 0.5 then 4, 5, 7.5, and 10). Then, we calculated $AIC_c$ for each model and selected the one with the lowest value conditioned on it having fewer or the same number of coefficients as presence sites (Warren & Siefert 2011). In preliminary analyses we tried turning threshold and hinge features off to create smoother models, but found that Maxent had trouble recreating response curves for species that responded with a logistic form to the TRUE variable. As a result, we used all possible features. Maxent was trained using 10,000 randomly located background sites (Phillips et al. 2006). Maxent does not allow site-level weighting.

**Boosted regression trees (BRTs)**

BRTs are also potentially highly flexible models based on chaining a large number of classification and regression trees (CART) serially (Elith et al. 2008). Each CART divides the predictor space into a series of hyperrectangles based on divisions that best separate cases (presence from absence or presence from background sites). Successive CARTs are constructed on a randomly selected subset of the data (the "bag" fraction), which reduces model bias. We calibrated BRTs using the *trainBrt* function in the [redacted for double-blind peer review]package [redacted for double-blind peer review]. Key parameters for calibrating BRTs include the tree complexity (potential number of splits in any one CART), the learning rate (the weighting applied to each CART in the overall model), and the total number of CARTs used in the final model. *trainBrt* calls the *gbm.step* function in the *dismo* package (Hijmans et al. 2018) which finds the optimal number of trees by comparing the deviance across a series of models using an increasing number of final trees. The function *trainBrt* also calibrates tree depth and learning rate (and bag fraction, which we did not explore). On some occasions, BRTs did not converge with the given data set, combination of parameters, and random number seed. The *trainBrt* function attempts to address this by slightly altering values for tree complexity, learning rate, maximum number of trees, and step size then trying again up to 5 times with different values each time. Nevertheless, in certain circumstances we were unable to obtain models that converged for some (sometimes all) cases. We note this in the main text and indicate it in figures with bar plots by sizing bar width proportional to the total number of models that converged. We trained BRT models using 200 randomly located background sites (Barbet-Massin et al. 2012).

To expedite processing *Experiments 1* through *6* with one TRUE and one FALSE variable, we conducted a preliminary scan of tuning parameters using the scenario described in *Experiment 1* where a species responds in a logistic fashion to a TRUE variable with a linear gradient spanning the landscape, but the model is also presented data on a FALSE variable that has randomly allocated values and is thus uncorrelated with the distribution of the species or the TRUE variable. We simulated 100 iterations each with different instantiations of the FALSE variable and different sets of training and test sites. We then used the *trainBrt* function to conduct a grid search across all possible combinations learning rate (0.00005, 0.0001, 0.0005, 0.001, 0.005, 0.01), tree complexity (1, 2, 3, 6), and number of trees (from 1000 to 8000 in steps of 50). The bag fraction was set at 0.6. From this set of models, we then selected a smaller set of parameters for a restricted grid search when processing *Experiments 1* through *6*. Specifically, we used: 1.1 times the maximum number of optimal trees across all 100 models, rounded up to the nearest 50; the mean learning rate and the 25th and 75th quantile values of the mean learning rate; and the mean (rounded) tree complexity and the 25th and 75th quantile values of tree complexity. If the mean tree complexity was equal to 1 then we used it and the 75th quantile. If the 25th quantile of tree complexity was equal to the mean we used the mean minus 1 instead of the 25th quantile (unless the mean value was equal to 1). If the 75th quantile was equal to the mean we used the mean plus 1 in its stead. The bag fraction was kept at 0.6.

For *Experiments 7* through *9* we trained models using a restricted set of parameters somewhat broader than the restricted set selected in the first tuning exercise but also narrower than the full set of parameters evaluated in that exercise. Specifically, we tuned models using a grid search of learning rate (0.005, 0.001, 0.0005, 0.0001), tree complexity (1, 2, 3), maximum trees (from 1000 to 8000 trees in steps of 50), and a bag fraction of 0.6.

[redacted for double-blind peer review]

[redacted for double-blind peer review]

Warren, D.L. and S.N. Siefert. 2011. Ecological niche modeling in Maxent: The importance of model complexity and the performance of model selection criteria. Ecological Applications 21:335-342.

Wood, S.N. 2006. Generalized Additive Models: An Introduction with R. Chapman and Hall/CRC, Boca Raton. 392 pp.
