## Appendix 2 for "Testing the ability of species distribution models to infer variable importance"

### Maps of landscapes and species

[Author names withheld for double-blind peer review.]

This appendix presents maps of the landscape and species for each of the experiments described in the main text. *Experiments 1* through *5* use a square landscape with one TRUE (influential) variable with a linear gradient and one FALSE (uninfluential) variable with a spatially random distribution. Unless otherwise stated, each variable ranged from -1 to 1 and landscaped were 104 cells on a side. The species' maps represent the probability of occurrence. Training and test occurrences and test absences were drawn from the species' maps using a Bernoulli trial with the probability of success equal to the probability of occurrence.

*Experiment 6* uses a circular landscape in which the FALSE variable is rotated relative to the TRUE variable to change the degree of collinearity (correlation) between them. We used a circular landscape to keep the univariate frequency distribution of each variable unchanged as one was rotated relative to the other.

*Experiment 7* also uses a circular landscape, but with two TRUE variables. The variables can be rotated relative to one another, and they can interact to define the species' niche (Eq. 3).

All landscapes were generated using the genesis function in the [redacted for double-blind peer review] package ver. 0.5.0 [redacted for double-blind peer review] for R (R Core Team 2020).

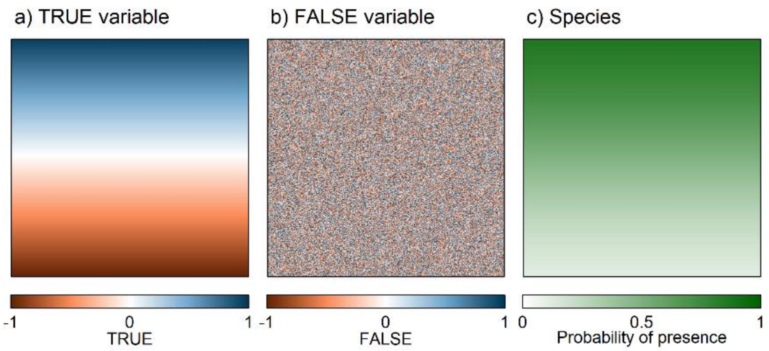


**Figure S1 (*Experiment 1: Simple scenario* and *Experiment 2: Sample size*)**. Spatial arrangement of landscape and species. (a) The "TRUE" variable has a linear trend across the landscape with a range of (-1, 1). (b) The "FALSE" variable has randomly distributed values, also in the range (-1, 1). (c) The species responds in a logistic fashion to the TRUE variable. Training and test occurrences are sampled from the species' distribution proportional to cells' probability of occurrence (the "species" map). The landscape is 1024 cells on a side. Scales along the bottom display the range of the values shown in the maps.

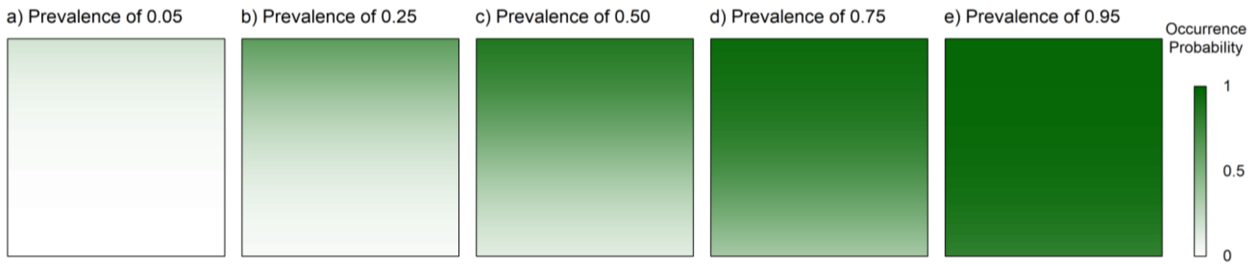


**Figure S2 (*Experiment 3: Prevalence*)**. Prevalence (mean probability of occurrence) was varied from 0.05 to 0.95 along a series of 9 steps (only 5 are shown here) by "shifting" the species' distribution with an offset parameter (Eq. 2). The distribution of the TRUE and FALSE variables remained unchanged and are as in Fig. S1. The landscape is 1024 cells on a side.

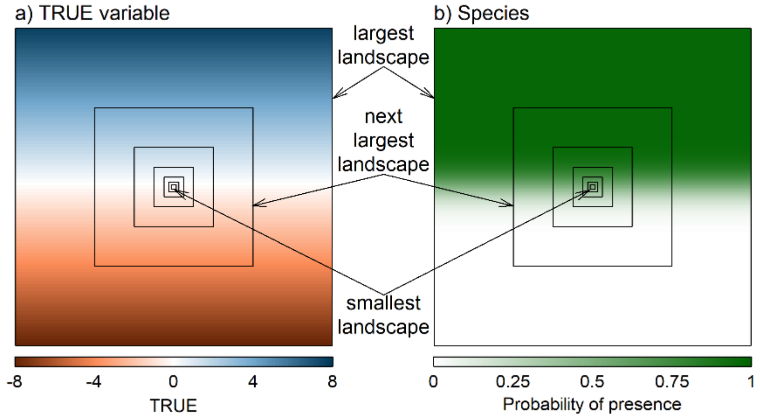


**Figure S3 (*Experiment 4: Extent*)**. Study region extent was increased while keeping the species' prevalence unchanged. To test the effects of the relationship between extent and the range of environmental variables on assessing variable importance, we simulated seven landscapes of increasing size (128 cells on a side to 8192 cells on a side) with concomitant increases in the overall range of the TRUE variable (from 0.25 at the smallest extent to 16 at the largest). The species had a logistic response to the TRUE variable. Increasing extent increased the range of the probability of presence across the landscape. Prevalence (mean suitability) of the species was kept constant across the landscapes.

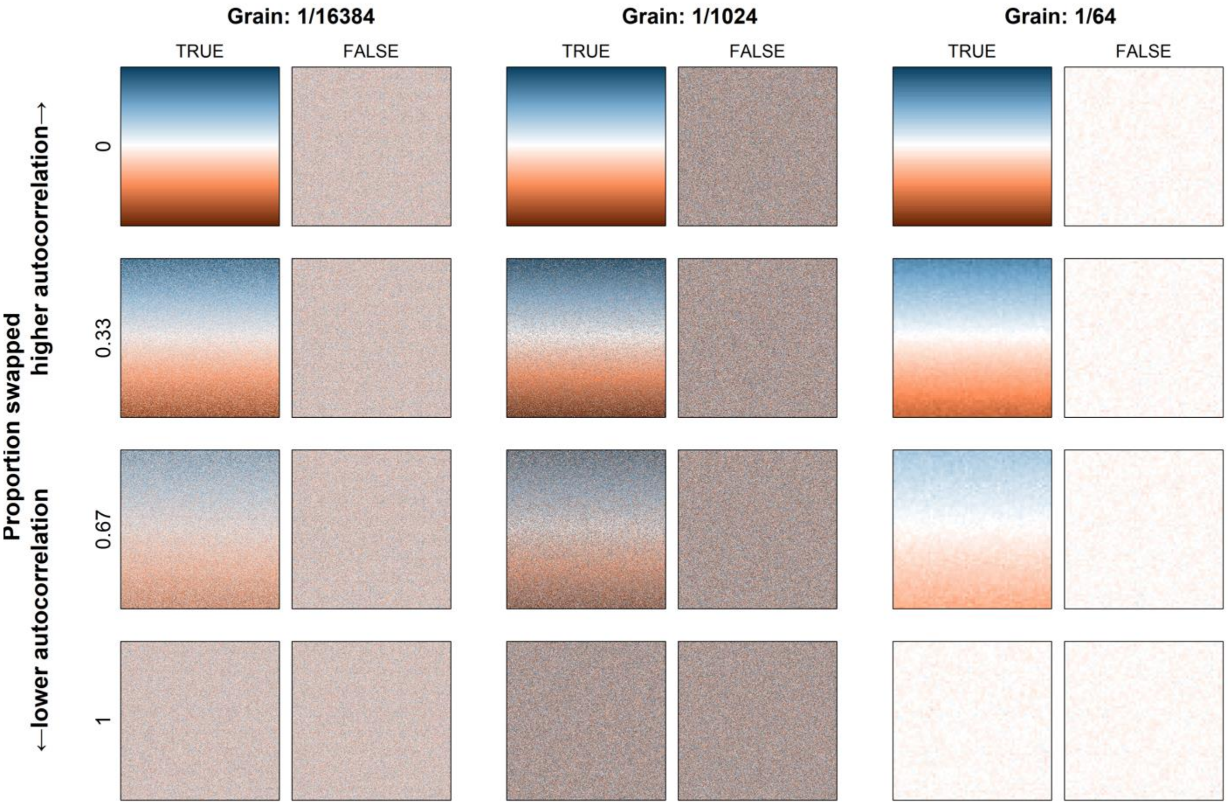


**Figure S4 (*Experiment 5: Spatial resolution and autocorrelation of environmental data*).** Landscapes for testing the effects of grain size (spatial resolution) of data and spatial autocorrelation. The species was assumed to respond to the environment at a "native" grain size of 1/1024$^{th}$ of the linear dimension of the landscape. The environmental data was resampled to a resolution of 1/16,384th (smaller cells) or 1/64th (large cells) of the landscape's linear dimension. Spatial autocorrelation in the environment mediates the effect of spatial resolution, so we varied autocorrelation for each variable by randomly swapping cell values between none, 1/3, 2/3, and all of the cells. Increasing the proportion of cells swapped reduces spatial autocorrelation in the TRUE variable because cells of similar value are less likely to be close to one another. Each TRUE was paired with the FALSE with the same level of spatial autocorrelation and grain (e.g., the TRUE in the upper left was paired with the FALSE to the immediate right of it). Values range from -1 (brown) to 1 (blue).

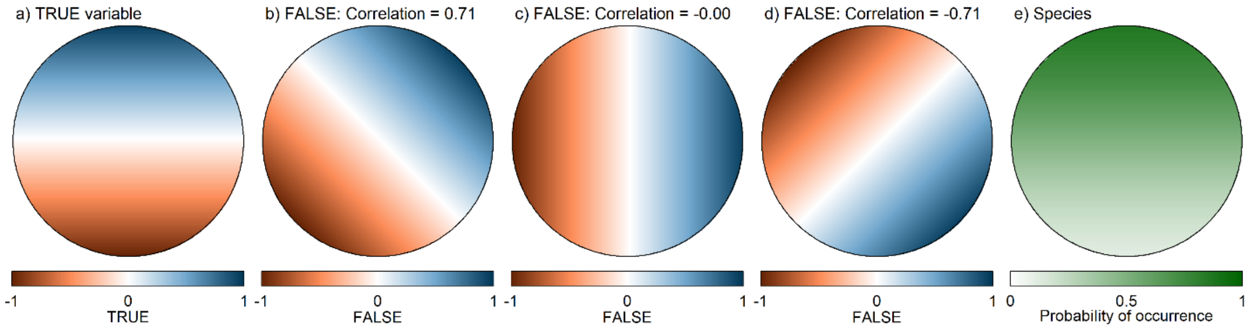


**Figure S5 (*Experiment 6: Collinearity*)**. Collinearity between environmental variables was varied by rotating the FALSE variable in relation to geographic the orientation of the TRUE variable on a circular landscape. In this scenario we created a circular landscape with a TRUE variable (a) that influenced the species' distribution. The model was also presented with data on a FALSE variable (b through d) that did not influence distribution. The FALSE variable was rotated relative to the gradient of the TRUE variable from 22.5° to 157.5° in steps of 22.5°. The rotations changed the correlation with TRUE from 0.92 to -0.92. Rotations of 45° (b), 90° (c), and 135° (d) are shown here. Since the species only responds to the TRUE variable, its distribution remains constant regardless of the rotation of FALSE relative to TRUE (e). However, the SDM algorithm is provided information on both TRUE and FALSE.

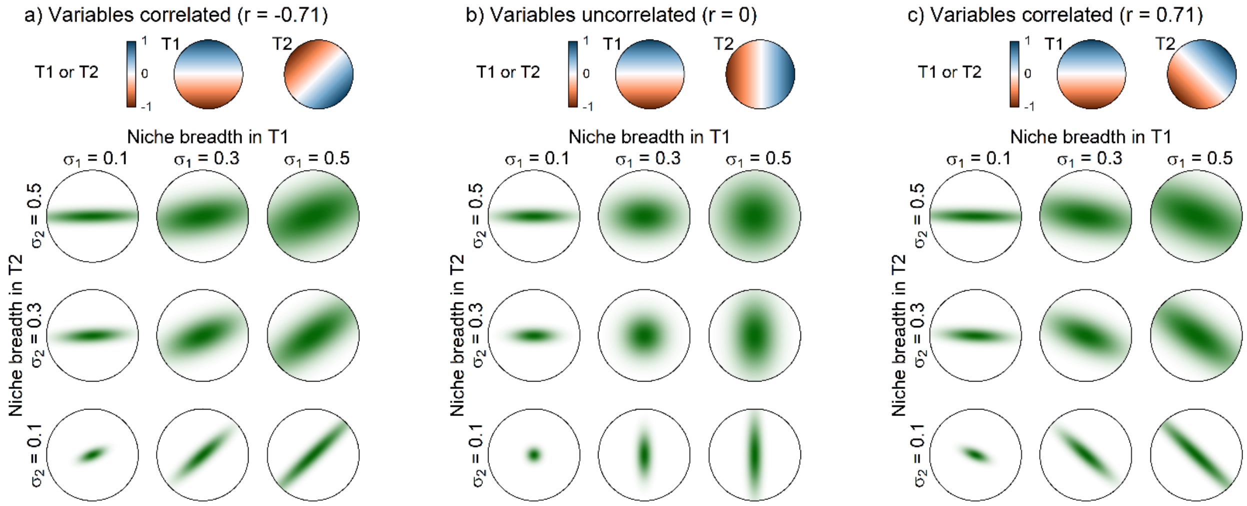


**Figure S6 (*Experiment 7*)**. Two variables act independently to shape the species' niche and distribution. The top row in each panel displays the distribution of two TRUE variables (T1 and T2) on the landscape. The bottom three rows show the species' distribution on the landscapes as a function of niche breadth in T1 and T2. Larger values of $\sigma_i$ connote broader niche breadth and thus less restriction by the respective variable. As a result, reducing $\sigma_i$ causes the variable to restrict its distribution more (i.e., the variable is more important). In *Experiment 7* we varied niche breadth while using a 90° rotation between T1 and T2 so they were uncorrelated (panel b). In *Experiment 8* we varied the correlation between T1 and T2 from -0.71 to 0 to 0.72 by rotating T2 relative to T1 on the landscape (panels a and c). The range of values for the species' maps is (0, 1), with darker/greener shades indicating greater probabilities of occurrence.

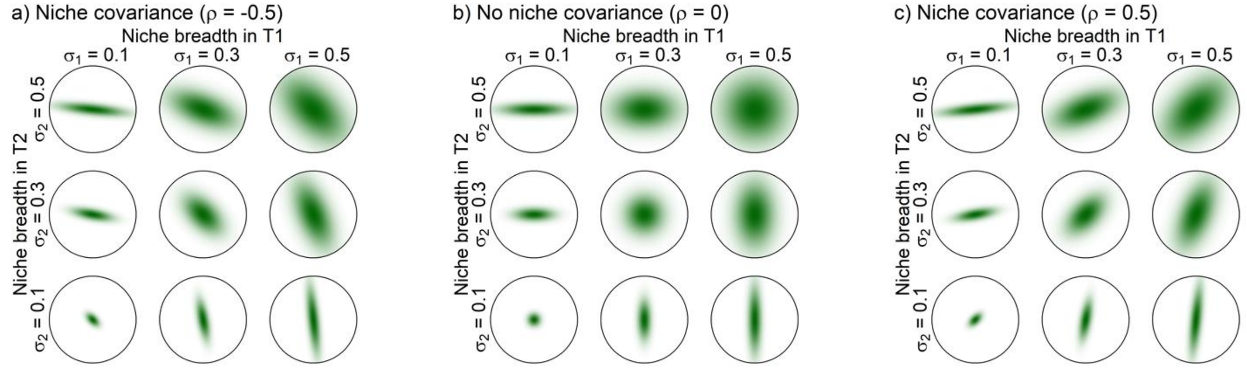


**Figure S7 (*Experiment 7*)**. Two TRUE variables (T1 and T2) interact to define the niche. In all cases shown here T1 and T2 were distributed orthogonally to one another on the landscape so were uncorrelated even though they interact to shape the niche (compare to Fig. 12a and c). Parameter $\rho$ determines the interaction between T1 and T2. The degree of limitation imposed by T1 and T2 are determined jointly by $\sigma_1$ and $\sigma_2$ plus the magnitude and sign of $\rho$. Panel (b) is the same as Fig. 12b but repeated here to aid visual comparison. The predicted probability of presence ranges from 0 (white) to 1 (dark green).
