## Appendix 3 for "Testing the ability of species distribution models to infer variable importance"

**Appendix 3: Summary of results for CBI, AUC, COR, and algorithm-specific tests**

[Author names withheld for double-blind peer review.]

### Overview of methods

In the main text of the article we present results for Maxent only using the Continuous Boyce Index (CBI) as test statistic for the permute-after-calibration test of variable importance. In this supplement we summarize results for all experiments using a TRUE and a FALSE variable for all modeling algorithms (GAMs, Maxent, and BRTs) and test statistics (CBI, $AUC_{pa}$—AUC calculated with presences and absences, $AUC_{bg}$—AUC calculated with presences and background sites, $COR_{pa}$—the correlation between permuted and unpermuted predictions at test presences and absence sites, and $COR_{bg}$—the correlation between permuted and unpermuted predictions at test presence and background sites). Asterisks (*) indicate test reliably discriminated between TRUE and FALSE.  Daggers (†) indicate the results were well-calibrated with the OMNI model. Note that in a few cases not all models converged (GAMs and BRTs) or yielded more than intercept-only models (Maxent). As a result, successful discrimination may be based on fewer than 100 models.

**Table S1. Sample size**

| **a) CBI** | Number of calibration presences | | | | | | | |
|---|---|---|---|---|---|---|---|---|
| Algorithm | 8 | 16 | 32 | 64 | 128 | 256 | 512 | 1024 |
| OMNI | * | * | * | * | * | * | * | * |
| GAM | * | * | * | * | * | * | * | * |
| Maxent | | | * | * | * | * | * | * |
| BRT | | | | * | * | * | * | * |

| **b) $AUC_{pa}$** | Number of calibration presences | | | | | | | |
|---|---|---|---|---|---|---|---|---|
| Algorithm | 8 | 16 | 32 | 64 | 128 | 256 | 512 | 1024 |
| OMNI | * | * | * | * | * | * | * | * |
| GAM | | | | | *† | *† | *† | *† |
| Maxent | | | * | * | * | *† | *† | *† |
| BRT | | | * | * | * | * | * | * |

| **c) $AUC_{bg}$** | Number of calibration presences | | | | | | | |
|---|---|---|---|---|---|---|---|---|
| Algorithm | 8 | 16 | 32 | 64 | 128 | 256 | 512 | 1024 |
| OMNI | * | * | * | * | * | * | * | * |
| GAM | | | | | * | *† | *† | *† |
| Maxent | | | * | * | * | * | * | * |
| BRT | | | * | * | * | * | * | * |

| **d) $COR_{pa}$** | Number of calibration presences | | | | | | | |
|---|---|---|---|---|---|---|---|---|
| Algorithm | 8 | 16 | 32 | 64 | 128 | 256 | 512 | 1024 |
| OMNI | * | * | * | * | * | * | * | * |
| GAM | * | * | *† | * | * | * | * | * |
| Maxent | | | | * | * | * | * | * |
| BRT | | | | * | * | * | * | * |

| **e) $COR_{bg}$** | Number of calibration presences | | | | | | | |
|---|---|---|---|---|---|---|---|---|
| Algorithm | 8 | 16 | 32 | 64 | 128 | 256 | 512 | 1024 |
| OMNI | * | * | * | * | * | * | * | * |
| GAM | * | * | * | * | * | * | * | * |
| Maxent | | | | * | * | * | * | * |
| BRT | | | | * | * | * | * | * |

| **f) Specific** | Number of calibration presences | | | | | | | |
|---|---|---|---|---|---|---|---|---|
| Algorithm | 8 | 16 | 32 | 64 | 128 | 256 | 512 | 1024 |
| Maxent Contrib. | | | | * | * | * | * | * |
| Maxent Permut. | | | | * | * | * | * | * |
| Maxent Gain | | | | | * | * | * | * |
| BRT | | | | | * | * | * | * |

**Table S2. Prevalence**

**a) CBI**

| Algorithm | 0.05 | 0.15 | 0.25 | 0.38 | 0.5 | 0.62 | 0.75 | 0.85 | 0.95 |
|---|---|---|---|---|---|---|---|---|---|
| OMNI | * | * | * | * | * | * | * | | |
| GAM | * | * | * | * | * | * | * | | |
| Maxent | * | * | * | * | * | * | * | | |
| BRT | * | * | * | * | * | * | | | |

(Columns: Prevalence)

**b) $AUC_{pa}$**

| Algorithm | 0.05 | 0.15 | 0.25 | 0.38 | 0.5 | 0.62 | 0.75 | 0.85 | 0.95 |
|---|---|---|---|---|---|---|---|---|---|
| OMNI | * | * | * | * | * | * | * | * | * |
| GAM | *† | *† | *† | * | * | * | | | |
| Maxent | *† | *† | *† | * | * | * | * | | |
| BRT | * | * | * | * | * | * | * | | |

(Columns: Prevalence)

**c) $AUC_{bg}$**

| Algorithm | 0.05 | 0.15 | 0.25 | 0.38 | 0.5 | 0.62 | 0.75 | 0.85 | 0.95 |
|---|---|---|---|---|---|---|---|---|---|
| OMNI | * | * | * | * | * | * | * | | |
| GAM | *† | *† | * | *† | * | *† | | | |
| Maxent | *† | * | * | * | * | * | * | | |
| BRT | * | * | * | * | * | * | * | | |

(Columns: Prevalence)

**d) $COR_{pa}$**

| Algorithm | 0.05 | 0.15 | 0.25 | 0.38 | 0.5 | 0.62 | 0.75 | 0.85 | 0.95 |
|---|---|---|---|---|---|---|---|---|---|
| OMNI | * | * | * | * | * | * | * | * | * |
| GAM | * | * | * | * | * | * | * | | |
| Maxent | * | * | * | * | * | * | | | |
| BRT | * | * | * | * | * | * | | | |

(Columns: Prevalence)

**e) $COR_{bg}$**

| Algorithm | 0.05 | 0.15 | 0.25 | 0.38 | 0.5 | 0.62 | 0.75 | 0.85 | 0.95 |
|---|---|---|---|---|---|---|---|---|---|
| OMNI | * | * | * | * | * | * | * | * | * |
| GAM | * | * | * | * | * | * | * | | |
| Maxent | * | * | * | * | * | * | | | |
| BRT | * | * | * | * | * | * | | | |

(Columns: Prevalence)

**f) Specific**

| Algorithm | 0.05 | 0.15 | 0.25 | 0.38 | 0.5 | 0.62 | 0.75 | 0.85 | 0.95 |
|---|---|---|---|---|---|---|---|---|---|
| Maxent Contrib. | * | * | * | * | * | * | | | |
| Maxent Permut. | * | * | * | * | * | * | | | |
| Maxent Gain | * | * | * | * | * | * | | | |
| BRT | * | * | * | * | * | | | | |

(Columns: Prevalence)

**Table S3. Study region extent**

| **a) CBI** | Study region extent (range of TRUE) | | | | | | |
|---|---|---|---|---|---|---|---|
| Algorithm | 0.25 | 0.5 | 1 | 2 | 4 | 8 | 16 |
| OMNI | | | * | * | * | * | * |
| GAM | | | * | * | * | * | * |
| Maxent | | | | * | * | * | * |
| BRT | | | | * | * | * | * |

| **b) $AUC_{pa}$** | Study region extent (range of TRUE) | | | | | | |
|---|---|---|---|---|---|---|---|
| Algorithm | 0.25 | 0.5 | 1 | 2 | 4 | 8 | 16 |
| OMNI | | * | * | * | * | * | * |
| GAM | | | | * | * | * | * |
| Maxent | | | * | * | * | * | * |
| BRT | | | * | * | * | * | * |

| **c) $AUC_{bg}$** | Study region extent (range of TRUE) | | | | | | |
|---|---|---|---|---|---|---|---|
| Algorithm | 0.25 | 0.5 | 1 | 2 | 4 | 8 | 16 |
| OMNI | | | * | * | * | * | * |
| GAM | | | | *† | * | *† | * |
| Maxent | | | * | * | * | * | * |
| BRT | | | | * | * | * | * |

| **d) $COR_{pa}$** | Study region extent (range of TRUE) | | | | | | |
|---|---|---|---|---|---|---|---|
| Algorithm | 0.25 | 0.5 | 1 | 2 | 4 | 8 | 16 |
| OMNI | * | * | * | * | * | * | * |
| GAM | * | | * | * | * | * | * |
| Maxent | | | * | * | * | * | * |
| BRT | | | | * | * | * | * |

| **e) $COR_{bg}$** | Study region extent (range of TRUE) | | | | | | |
|---|---|---|---|---|---|---|---|
| Algorithm | 0.25 | 0.5 | 1 | 2 | 4 | 8 | 16 |
| OMNI | * | * | * | * | * | * | * |
| GAM | * | | * | * | * | * | * |
| Maxent | | | * | * | * | * | * |
| BRT | | | | * | * | * | * |

| **f) Specific** | Study region extent (range of TRUE) | | | | | | |
|---|---|---|---|---|---|---|---|
| Algorithm | 0.25 | 0.5 | 1 | 2 | 4 | 8 | 16 |
| Maxent Contrib. | | | * | * | * | * | * |
| Maxent Permut. | | | * | * | * | * | * |
| Maxent Gain | | | | * | * | * | * |
| BRT | | | | * | * | * | * |

**Table S4. Environmental data resolution and spatial autocorrelation**

In each cell "o" indicates reliable discrimination by OMNI, "g" by GAMs, "m" by Maxent, and "b" by BRTs. A capitalized letter indicates reliable calibration vis-à-vis the omniscient model. For the algorithm-specific metrics all Maxent-based tests had the same discrimination capacity.

**a) CBI**

| | Grain size | | |
|---|---|---|---|
| Swapped | 1/16,384 | 1/1024 | 1/64 |
| 0 | ogmb | ogmb | ogmb |
| 1/3 | ogmb | ogmb | ogmb |
| 2/3 | ogmb | ogmb | |
| 1 | ogmb | ogmb | |

**b) $AUC_{pa}$**

| | Grain size | | |
|---|---|---|---|
| Swapped | 1/16,384 | 1/1024 | 1/64 |
| 0 | oGmb | ogmb | ogmb |
| 1/3 | oGmb | ogmb | oGMb |
| 2/3 | ogmb | oGmb | |
| 1 | oGmb | ogmb | |

**c) $AUC_{bg}$**

| | Grain size | | |
|---|---|---|---|
| Swapped | 1/16,384 | 1/1024 | 1/64 |
| 0 | ogmb | ogmb | oGmb |
| 1/3 | oGmb | oGmb | oGmb |
| 2/3 | ogmb | oGmb | |
| 1 | ogmb | oGmb | |

**d) $COR_{pa}$**

| | Grain size | | |
|---|---|---|---|
| Swapped | 1/16,384 | 1/1024 | 1/64 |
| 0 | ogmb | ogmb | ogmb |
| 1/3 | ogmb | ogmb | ogm |
| 2/3 | ogmb | ogmb | og |
| 1 | ogmb | ogmb | |

**e) $COR_{bg}$**

| | Grain size | | |
|---|---|---|---|
| Swapped | 1/16,384 | 1/1024 | 1/64 |
| 0 | ogmb | ogmb | ogmb |
| 1/3 | ogmb | ogmb | ogm |
| 2/3 | ogmb | ogmb | og |
| 1 | ogmb | ogmb | |

**f) Specific**

| | Grain size | | |
|---|---|---|---|
| Swapped | 1/16,384 | 1/1024 | 1/64 |
| 0 | mb | mb | mb |
| 1/3 | mb | mb | |
| 2/3 | mb | mb | |
| 1 | mb | mb | |

**Table S5. Collinearity**

**a) CBI**

| Algorithm | Correlation between TRUE and FALSE | | | | | | |
|---|---|---|---|---|---|---|---|
| | -0.91 | -0.71 | -0.38 | 0 | 0.38 | -0.71 | -0.92 |
| OMNI | * | * | * | * | * | * | * |
| GAM | | * | * | * | * | * | |
| Maxent | | * | * | * | * | * | |
| BRT | | * | * | * | * | | |

**b) $AUC_{pa}$**

| Algorithm | Correlation between TRUE and FALSE | | | | | | |
|---|---|---|---|---|---|---|---|
| | -0.91 | -0.71 | -0.38 | 0 | 0.38 | -0.71 | -0.92 |
| OMNI | * | * | * | * | * | * | * |
| GAM | | * | *† | *† | * | * | |
| Maxent | | * | * | * | * | * | |
| BRT | | * | * | * | * | * | |

**c) $AUC_{bg}$**

| Algorithm | Correlation between TRUE and FALSE | | | | | | |
|---|---|---|---|---|---|---|---|
| | -0.91 | -0.71 | -0.38 | 0 | 0.38 | -0.71 | -0.92 |
| OMNI | * | * | * | * | * | * | * |
| GAM | | * | * | * | *† | * | |
| Maxent | | * | * | * | * | * | |
| BRT | | * | * | * | * | * | |

**d) $COR_{pa}$**

| Algorithm | Correlation between TRUE and FALSE | | | | | | |
|---|---|---|---|---|---|---|---|
| | -0.91 | -0.71 | -0.38 | 0 | 0.38 | -0.71 | -0.92 |
| OMNI | * | * | * | * | * | * | * |
| GAM | | * | * | * | * | * | |
| Maxent | | * | * | * | * | * | |
| BRT | | * | * | * | * | * | |

**e) $COR_{bg}$**

| Algorithm | Correlation between TRUE and FALSE | | | | | | |
|---|---|---|---|---|---|---|---|
| | -0.91 | -0.71 | -0.38 | 0 | 0.38 | -0.71 | -0.92 |
| OMNI | * | * | * | * | * | * | * |
| GAM | | * | * | * | * | * | |
| Maxent | | * | * | * | * | * | |
| BRT | | * | * | * | * | * | |

**f) Specific**

| Algorithm | Correlation between TRUE and FALSE | | | | | | |
|---|---|---|---|---|---|---|---|
| | -0.91 | -0.71 | -0.38 | 0 | 0.38 | -0.71 | -0.92 |
| Maxent Contrib. | | * | * | * | * | * | |
| Maxent Permut. | | * | * | * | * | * | |
| Maxent Gain | | | * | * | * | | |
| BRT | | * | | * | * | * | |
