## Appendix 4 for "Testing the ability of species distribution models to infer variable importance"

**Appendix 4: Full results using CBI as a test statistic**

[Author names withheld for double-blind peer review.]

### Overview of methods

In the main text of the article we present results for Maxent only using the Continuous Boyce Index (CBI) as test statistic for the permute-after-calibration test of variable importance. In this supplement we present results for GAMs, Maxent, and BRTs using CBI as the test statistic. CBI indicates how well a model's output serves as an index of the actual probability of presence (Boyce et al. 2002; Hirzel et al. 2006). It is calculated by first binning predictions to background sites into overlapping bins (we used 1001 bins). The proportion of background sites in each bin represents the expected number of test presences that should occur in each bin based on predictions to each test site. The ratio of observed to expected frequency of test occurrences in each bin is then calculated. Finally, bins are ranked from least to greatest expected values and the Pearson rank correlation coefficient calculated between the rank and the observed-to-expected ratio. Values of CBI >0 indicate the model is better than a random allocation of occurrences in proportion to expectation and values <0 indicate the model's predictions are inversely related to the expectation.

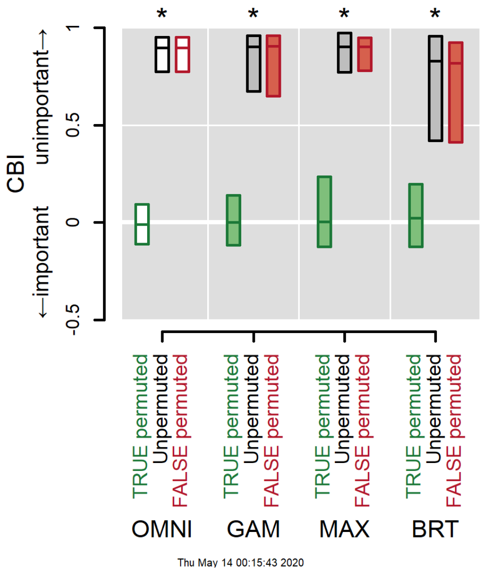


**Figure S1:** ***Experiment 1*****—Simple scenario**. A simple scenario with an influential TRUE variable and an uninfluential FALSE variable that was "mistakenly" presented to the model. All models successfully discriminated between the two variables (no overlap between green and red bars, indicated by asterisks). If a variable is important then permuting its values will lower model performance, here measured using CBI. The OMNI ("omniscient") model serves as a benchmark against which to assess the ability of the other models.

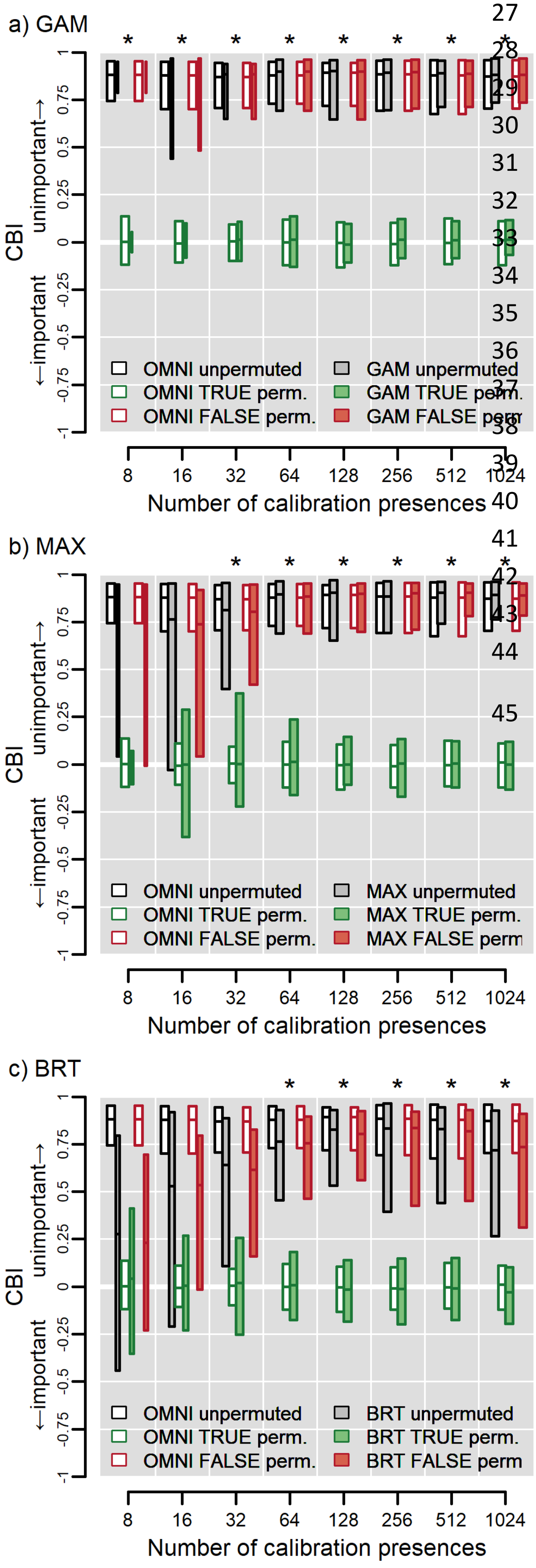


**Figure S2:** ***Experiment 2—Sample size***. GAMs can reliably discriminate between an influential and uninfluential variable at all training sample sizes, although some models did not converge when sample size was small (bar width is proportional to number of converged models). Maxent requires ≥32 and BRTs ≥64 occurrences for reliably discrimination. Asterisks indicate cases where both OMNI and the SDM successfully discriminate between TRUE and FALSE at the given number of presences. Bars represent the inner 95% of values of COR across 100 models on 100 data iterations of training and test sites at the given sample size. The horizontal line in each bar represents the median value. Results for OMNI are repeated across all panels. See Box 1 in the main text for further guidance on interpretation.

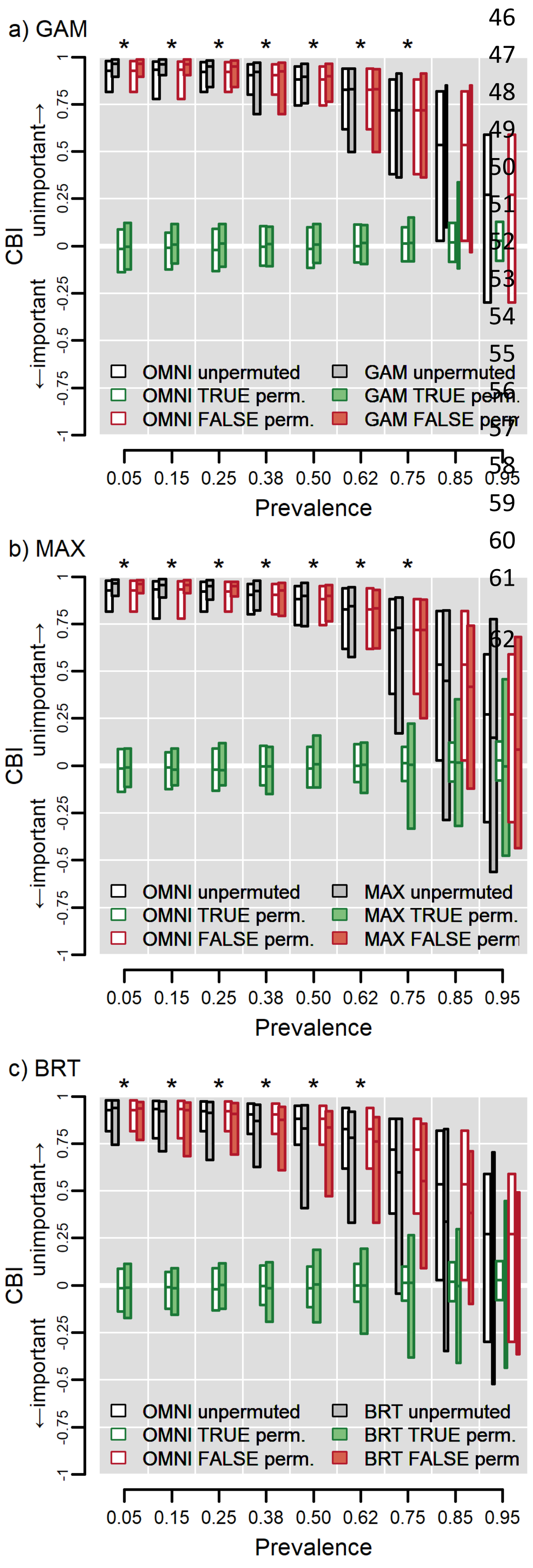


**Figure S3:** ***Experiment 3—Prevalence***. Models measure variable importance most accurately when a species occupies not more than ~62-75% of the study region. Asterisks indicate both OMNI and the SDM successfully discriminate between TRUE and FALSE at the given level of prevalence. Bars represent the inner 95% of values of $AUC_{pa}$ across 100 models for 100 data iterations of training and test sites at the given prevalence. The horizontal line in each bar represents the median value. Bar width is proportional to the number of iterations in which the respective SDM algorithm converged. Results for OMNI are repeated across all panels. See Box 1 for further guidance on interpretation.

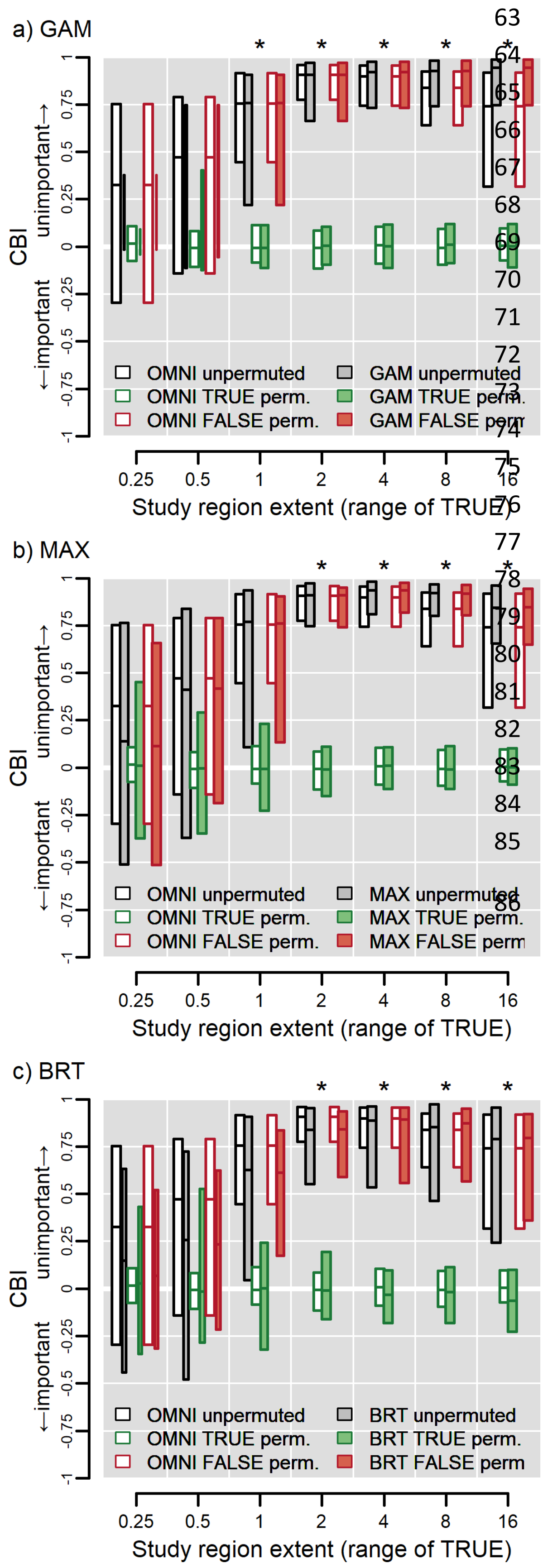


**Figure S4:** ***Experiment 4—Study region extent***. Models cannot reliably measure variable importance when study region extent is too small to encompass suitable variation in driving variables, even if the model is "omniscient." The x-axis in each panel displays the range of the TRUE variable (0. 25 for the smallest landscape and 16 for the largest) and thus reflects the effect of increasing the spatial extent of the study area (ranging 128 to 8192 cells on a side). Asterisks indicate OMNI and the SDM both successfully discriminate between TRUE and FALSE at the given extent (range of TRUE). Bars represent the inner 95% of values of $AUC_{pa}$ from 100 models for 100 data iterations of training and test sites at the given spatial extent. The horizontal line in each bar represents the median value. Bar width is proportional to the number of iterations in which the model successfully converged (<100 for GAMs and BRTs at ranges ≤1). Results for OMNI are repeated across all panels. See Box 1 for further guidance on interpretation.

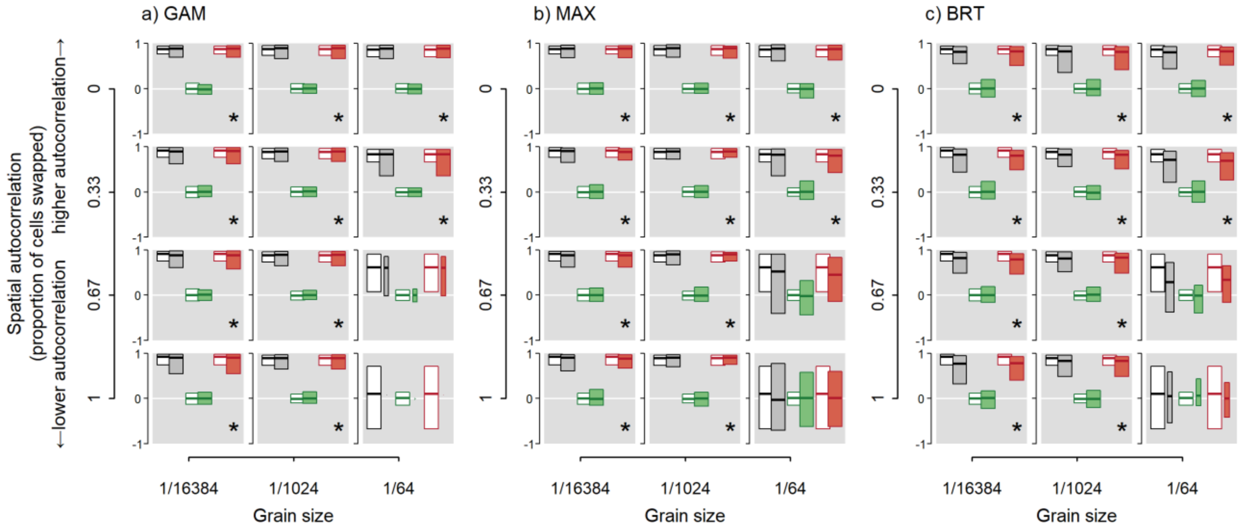


***Figure S5: Experiment 5—Spatial resolution and autocorrelation of environmental data*.** Models successfully discriminate between TRUE and FALSE variables regardless of spatial resolution of environmental data and spatial autocorrelation except when resolution is coarser than the perceptual scale of the species and autocorrelation is low. The resolution at which the species perceives the landscape is at 1/1024 (middle column in each panel). Bars in each subplot of each panel represent the inner 95% of values of CBI from 100 models for 100 data iterations of training and test sites at the given spatial resolution and magnitude of spatial autocorrelation. The horizontal line in each bar represents the median value. Bar width is proportional to the number of iterations in which the model successfully converged. Asterisks in the lower right corner of each subplot indicates both OMNI and the SDM successfully discriminate between the TRUE and FALSE variables. Results for OMNI are repeated across all panels. See Box 1 for further guidance on interpretation of the subplots in each panel.

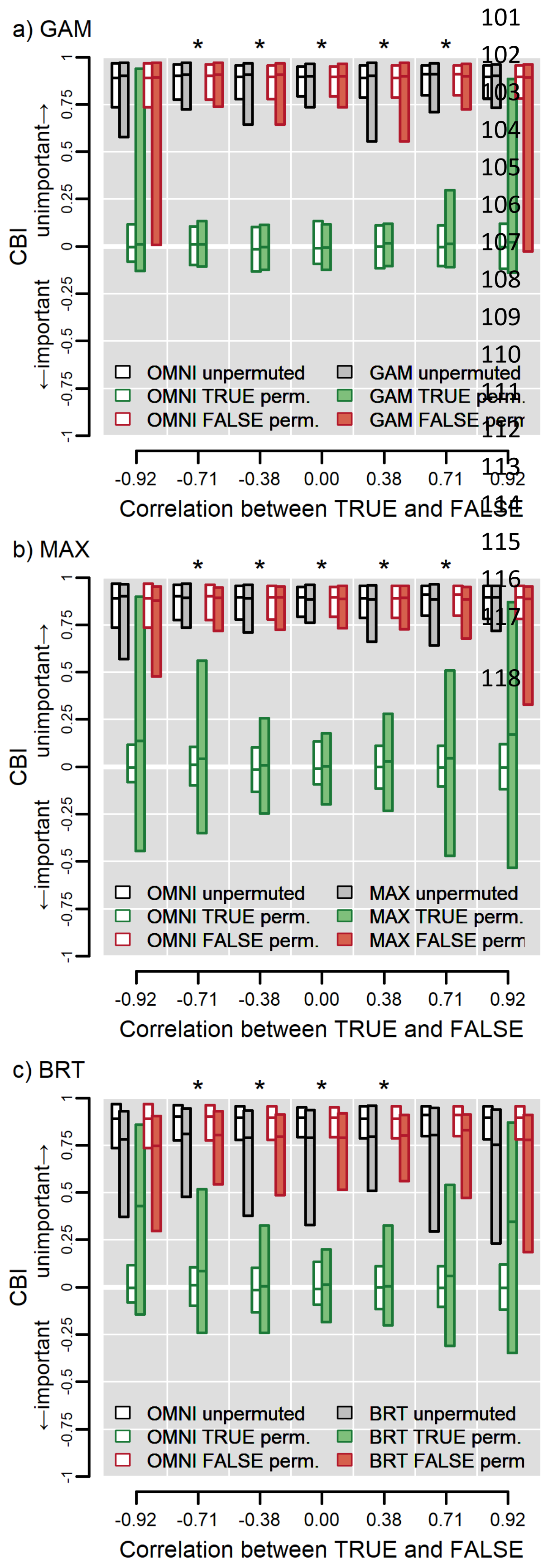


**Figure S6: *Experiment 6—Collinearity*.** Models have increasingly worse accuracy as the magnitude of correlation increases, especially when |*r*| >0.71. In each panel the ordinate reflects the correlation between the FALSE and TRUE variables which changes as the gradient in FALSE is rotated on the landscape relative to the gradient in TRUE. Asterisks indicates both OMNI and the SDM successfully discriminate between TRUE and FALSE at the given level of correlation. Bars represent the distribution of the inner 95% of values of $AUC_{pa}$ from 100 models for 100 data iterations of the species and landscape at the given level of correlation. Results from OMNI are repeated across panels. See Box 1 for further guidance on interpretation.

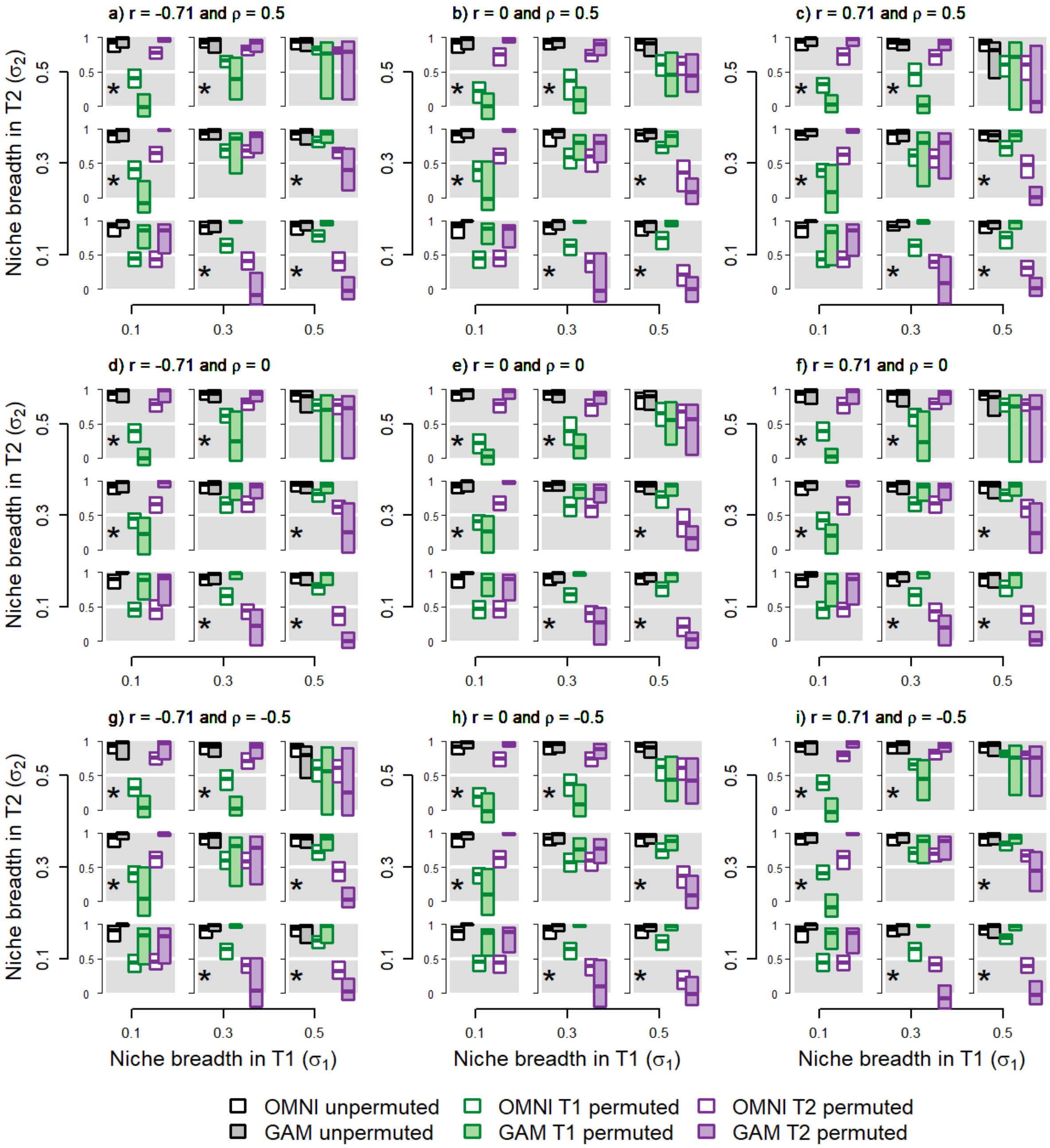


**Figure S7:** ***Experiments 7-9—Two influential variables modeled with GAMs***. Effect of niche breadth, collinearity, and niche covariance on inferences of variable importance using GAMs measured using CBI. Two variables, T1 and T2, each influence the niche according to the species' niche breadth in each. Each subpanel represents a landscape occupied by a species with the given niche breadths ($\sigma_1$ and $\sigma_2$), correlation between predictors ($r$) and niche covariance ($\rho$). Panel e corresponds to Fig. 4a in the main text (changing niche breadth with no correlation between predictors, no niche covariance). Asterisks indicate reliable discrimination and Latin crosses successful calibration. See Box 1 for further details on interpreting each subpanel in a panel.

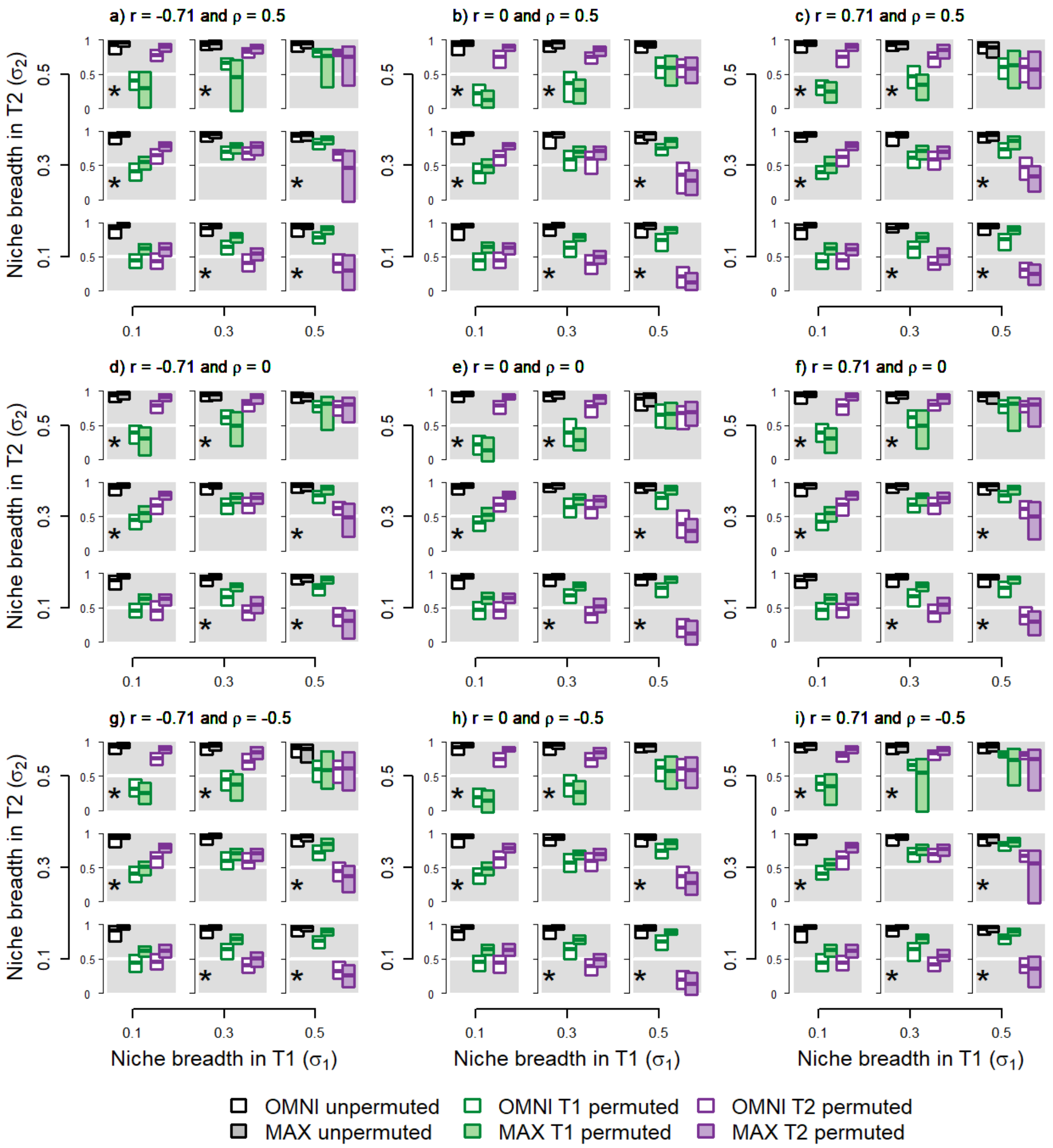


**Figure S8: *Experiments 7-9—Two influential variables modeled with Maxent*.** Effect of niche breadth, collinearity, and niche covariance on inferences of variable importance using Maxent measured using CBI. Two variables, T1 and T2, each influence the niche according to the species' niche breadth in each. Each subpanel represents a landscape occupied by a species with the given niche breadths ($\sigma_1$ and $\sigma_2$), correlation between predictors ($r$) and niche covariance ($\rho$). Panel e corresponds to Fig. 4a in the main text (changing niche breadth with no correlation between predictors, no niche covariance). Asterisks indicate reliable discrimination and Latin crosses successful calibration. See Box 1 for further details on interpreting each subpanel in a panel.

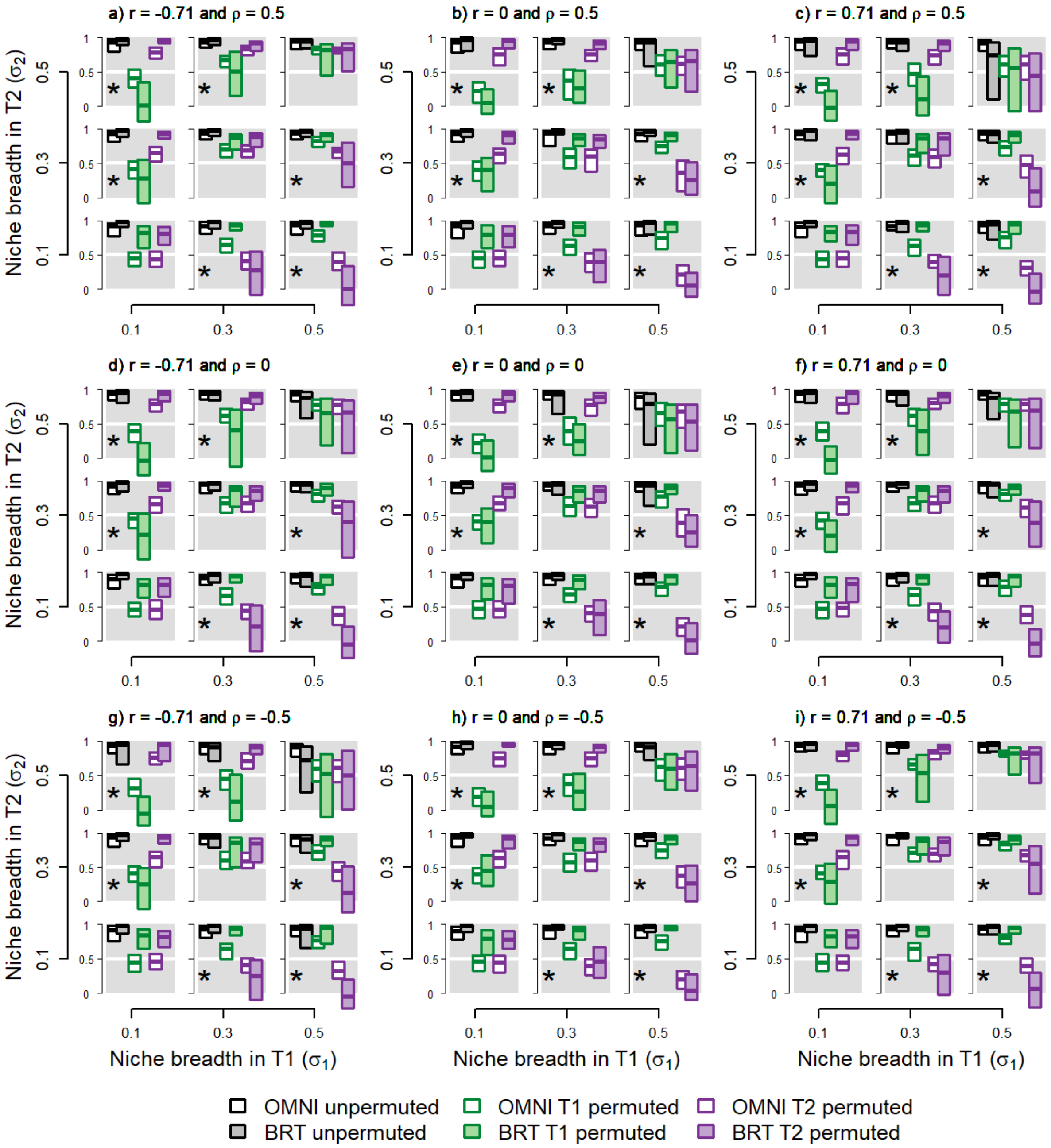


**Figure S9:** ***Experiments 7-9—Two influential variables modeled with BRTs***. Effect of niche breadth, collinearity, and niche covariance on inferences of variable importance using BRTs measured using CBI. Two variables, T1 and T2, each influence the niche according to the species' niche breadth in each. Each subpanel represents a landscape occupied by a species with the given niche breadths ($\sigma_1$ and $\sigma_2$), correlation between predictors ($r$) and niche covariance ($\rho$). Panel e corresponds to Fig. 4a in the main text (changing niche breadth with no correlation between predictors, no niche covariance). Asterisks indicate reliable discrimination and Latin crosses successful calibration. See Box 1 for further details on interpreting each subpanel in a panel.

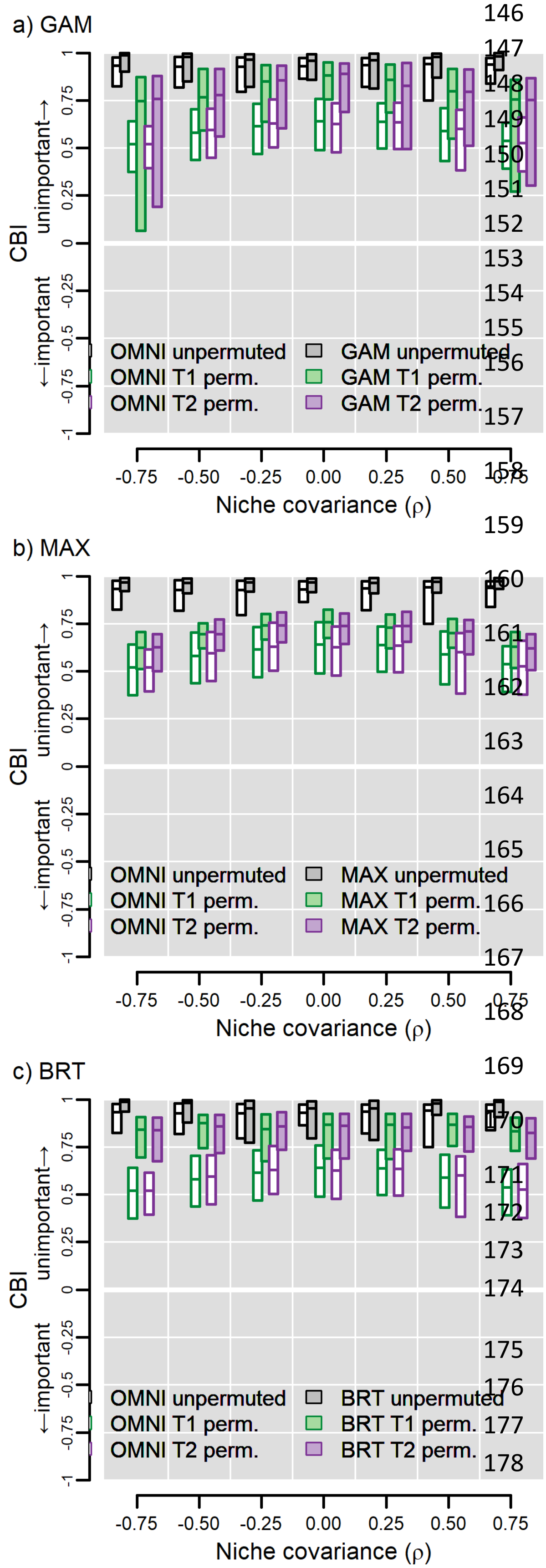


**Figure S10:** ***Experiment 8—Niche covariance*.** Variable importance increases as the magnitude of interaction between predictors increases. Here, niche width was held constant ($\sigma_1 = \sigma_2 = 0.3$) and the variables were uncorrelated on the landscape ($r = 0$). Since the variables had equal niche width, they should have the same influence. Asterisks indicate reliable discrimination and Latin crosses successful calibration. See Box 1 for further details on interpreting each subpanel in a panel.
