## Appendix 7 for "Testing the ability of species distribution models to infer variable importance"

**Appendix 7: Using the correlation test statistic calculated with presences and absences ($COR_{pa}$) as a test statistic for the permute-after-calibration test of variable importance**

[Author names withheld for double-blind peer review.]

### Overview of methods

In the main text of the article we present results using the Continuous Boyce Index (CBI) as test statistic for the permute-after-calibration test of variable importance. In this supplement we present figures complementary to those in the main text except we show results using the correlation between unpermuted and permuted predictions at test presence and absence sites ($COR_{pa}$; Breiman 2001). Unlike CBI or AUC, $COR_{pa}$ is not an indicator of a model's' predictive accuracy. As a result, $COR_{pa}$ can only be used to assess relative (versus absolute) importance of variables. Commonly used distribution and niche modeling software like Maxent (Phillips et al. 2006; Phillips & Dudík 2008) and the BIOMOD2 (Thuiller et al. 2009) and sdm (Naimi & Araújo 2016) packages for the R Statistical Environment (R Core Team 2018) can be used to estimate variable importance using AUC and/or COR.

In most cases the $COR_{pa}$ statistic for OMNI had almost no variation so appears as a horizontal line in boxplots. $COR_{pa}$ should equal 1 when the permuted and unpermuted predictions are exactly the same (the variable has no influence) and be <1 when the variable exerts influence. In practice the lowest expected value of $COR_{pa}$ should be ~0, but we found in some cases $COR_{pa}$ could fall below 0. We remind the reader that COR compares "unpermuted" and "permuted" model predictions. As a result, unlike figures for CBI and AUC, there are no results just for "unpermuted" predictions.

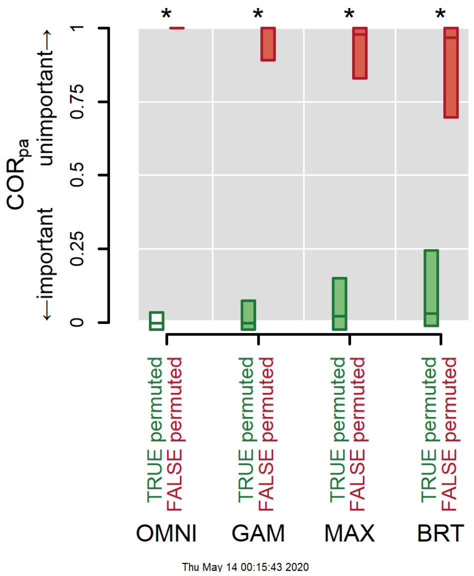


**Figure S1:** ***Experiment 1*****—Simple scenario**. A simple scenario with an influential TRUE variable and an uninfluential FALSE variable that was "mistakenly" presented to the model. All models successfully discriminated between the two variables (no overlap between green and red bars, indicated by asterisks). If a variable is important then permuting its values will lower model performance, here measured using $COR_{pa}$. The OMNI ("omniscient") model serves as a benchmark against which to assess the ability of the other models.

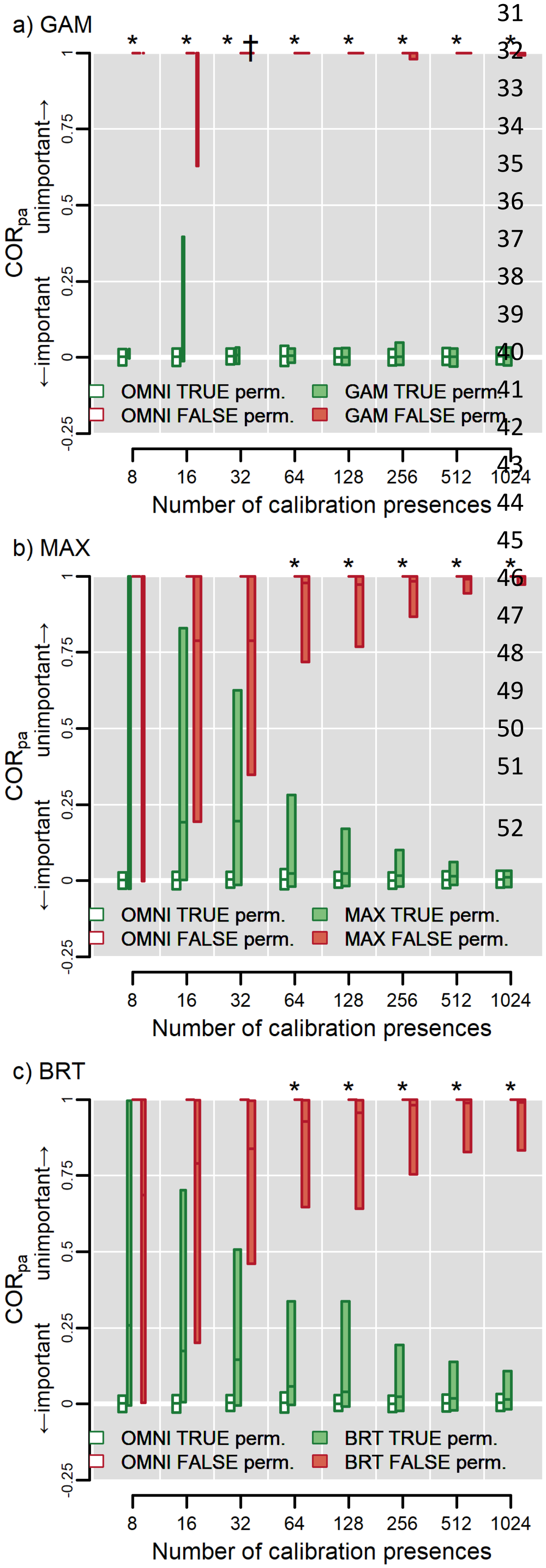


**Figure S2:** ***Experiment 2—Sample size***. Maxent and BRTs require ≥64 occurrences for reliable discrimination but GAMs can differentiate TRUE from FALSE at all sample sizes explored here. Bar width is proportional to the number of models that converged (GAMs and BRTs) or were more than intercept-only models (Maxent). Asterisks indicate cases where both OMNI and the SDM successfully discriminate between TRUE and FALSE at the given number of presences. Latin crosses indicate cases where the SDM is well-calibrated with OMNI (see main text, *General approach* under *Methods*). Bars represent the inner 95% of values of COR across 100 models on 100 data iterations of training and test sites at the given sample size. The horizontal line in each bar represents the median value. Results for OMNI are repeated across all panels. See Box 1 in the main text for further guidance on interpretation.

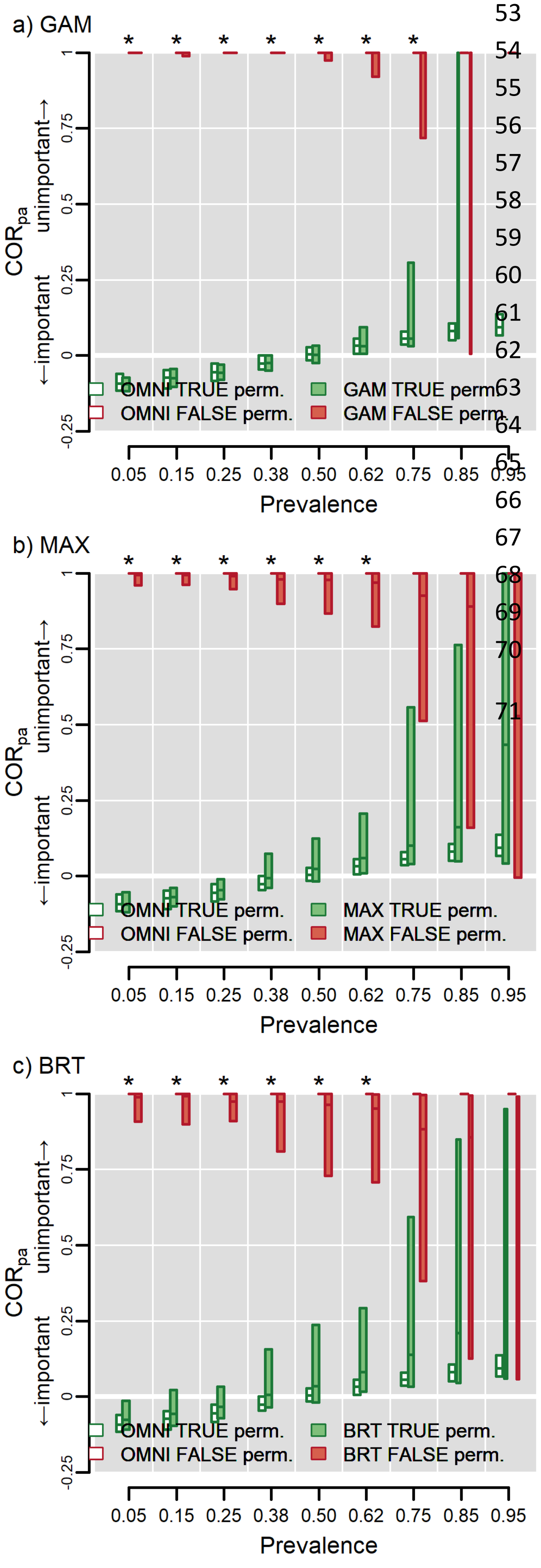


**Figure S3:** ***Experiment 3—Prevalence***. Models measure variable importance most accurately when a species occupies not more than ~62-75% of the study region. Asterisks indicate both OMNI and the SDM successfully discriminate between TRUE and FALSE at the given level of prevalence. Latin crosses indicate cases where the SDM is well-calibrated with OMNI (see main text, *General approach* under *Methods*). Bars represent the inner 95% of values of $COR_{pa}$ across 100 models for 100 data iterations of training and test sites at the given prevalence. The horizontal line in each bar represents the median value. Bar width is proportional to the number of iterations in which the respective SDM algorithm converged. Results for OMNI are repeated across all panels.  See Box 1 for further guidance on interpretation.

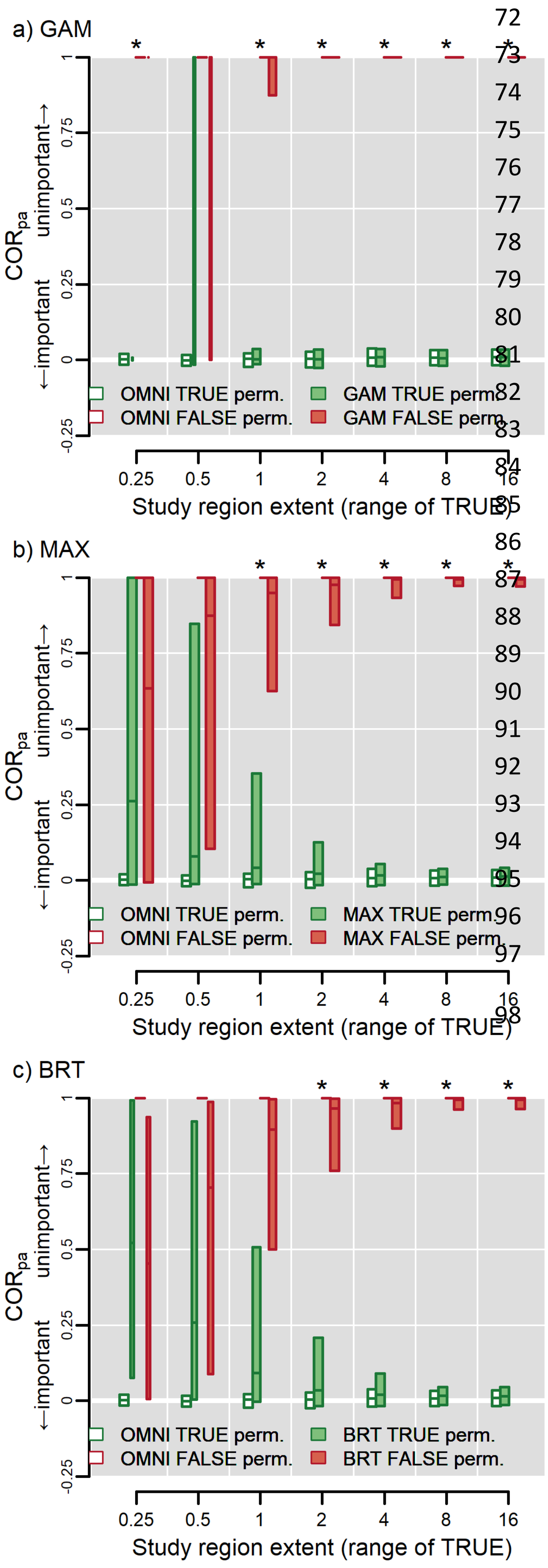


**Figure S4:** ***Experiment 4—Study region extent***. Models cannot reliably measure variable importance when study region extent is too small to encompass suitable variation in driving variables, even if the model is "omniscient." The x-axis in each panel displays the range of the TRUE variable (0. 25 for the smallest landscape and 16 for the largest) and thus reflects the effect of increasing the spatial extent of the study area (ranging 128 to 8192 cells on a side). Asterisks indicate OMNI and the SDM both successfully discriminate between TRUE and FALSE at the given extent (range of TRUE). Latin crosses indicate cases where the SDM is well-calibrated with OMNI (see main text, *General approach* under *Methods*). Bars represent the inner 95% of values of COR$_{pa}$ from 100 models for 100 data iterations of training and test sites at the given spatial extent. The horizontal line in each bar represents the median value. Bar width is proportional to the number of iterations in which the model successfully converged (<100 for GAMs and BRTs at ranges ≤1). Results for OMNI are repeated across all panels. See Box 1 for further guidance on interpretation.

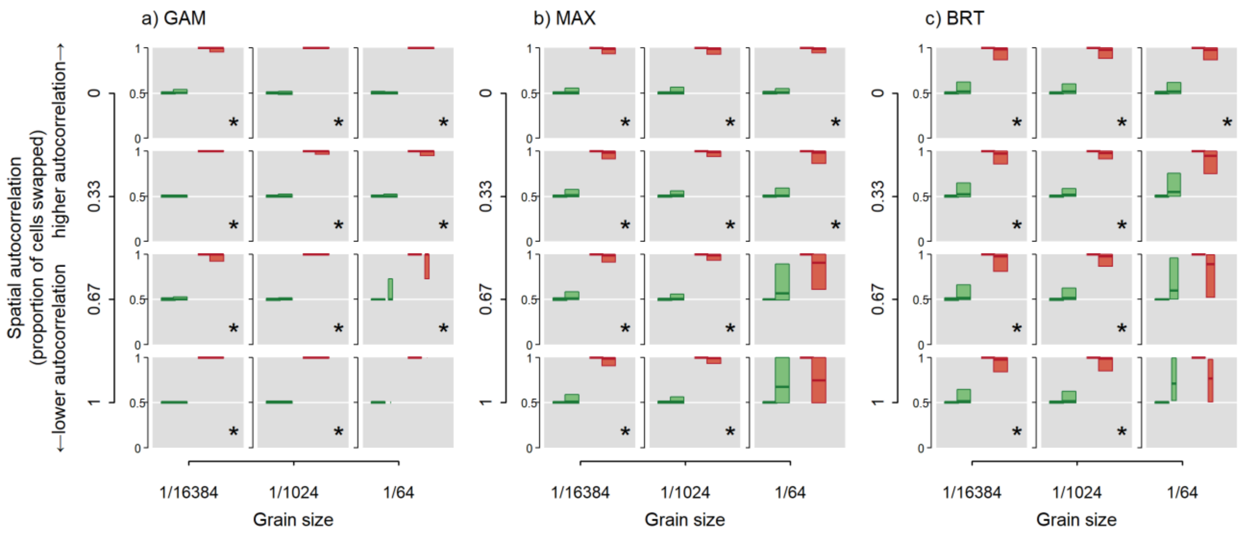


***Figure S5: Experiment 5—Spatial resolution and autocorrelation of environmental data***. Models successfully discriminate between TRUE and FALSE variables regardless of spatial resolution of environmental data and spatial autocorrelation except when resolution is coarser than the perceptual scale of the species and autocorrelation is low. The resolution at which the species perceives the landscape is at 1/1024 (middle column in each panel). Bars in each subplot of each panel represent the inner 95% of values of $COR_{pa}$ from 100 models for 100 data iterations of training and test sites at the given spatial resolution and magnitude of spatial autocorrelation. The horizontal line in each bar represents the median value. Bar width is proportional to the number of iterations in which the model successfully converged. Asterisks in the lower right corner of each subplot indicates both OMNI and the SDM successfully discriminate between the TRUE and FALSE variables. Latin crosses indicate cases where the SDM is well-calibrated with OMNI (see main text, *General approach* under *Methods*). Results for OMNI are repeated across all panels. See Box 1 for further guidance on interpretation of the subplots in each panel.

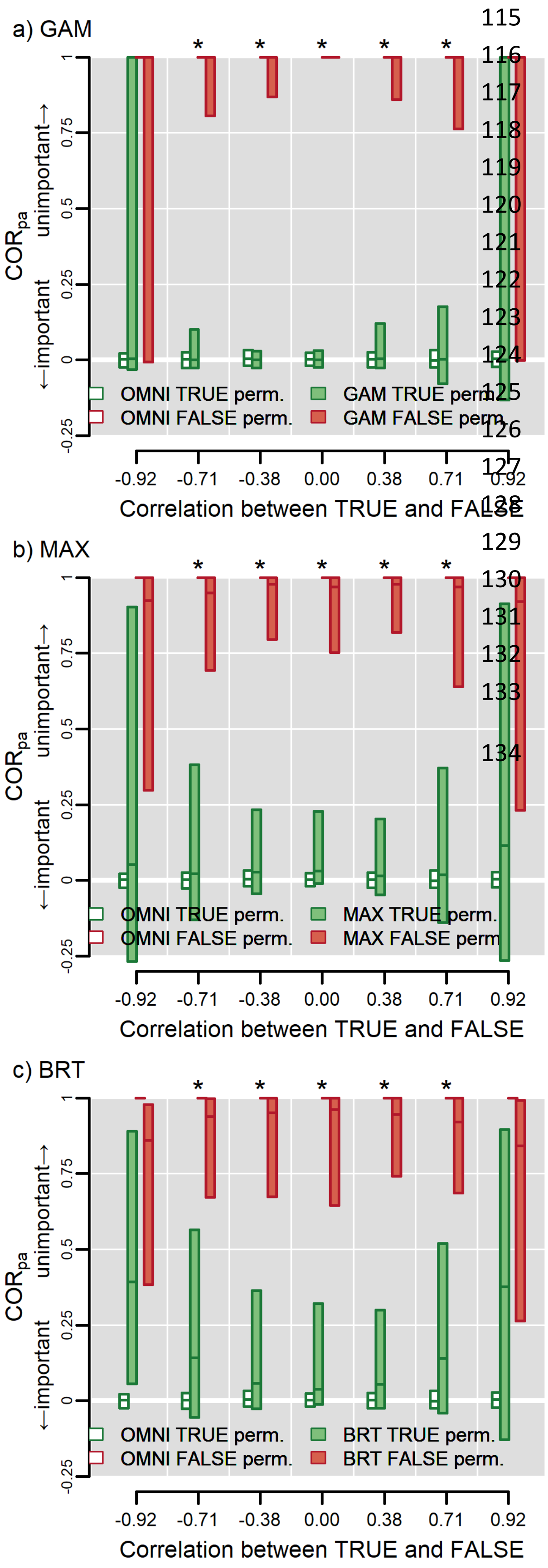


**Figure S6:** ***Experiment 6—Collinearity***. Models have increasingly worse accuracy as the magnitude of correlation increases, especially when |*r*| >0.71. In each panel the ordinate reflects the correlation between the FALSE and TRUE variables which changes as the gradient in FALSE is rotated on the landscape relative to the gradient in TRUE. Asterisks indicates both OMNI and the SDM successfully discriminate between TRUE and FALSE at the given level of correlation. Latin crosses indicate cases where the SDM is well-calibrated with OMNI (see main text, *General approach* under *Methods*). Bars represent the distribution of the inner 95% of values of $COR_{pa}$ from 100 models for 100 data iterations of the species and landscape at the given level of correlation. Results from OMNI are repeated across panels. See Box 1 for further guidance on interpretation.

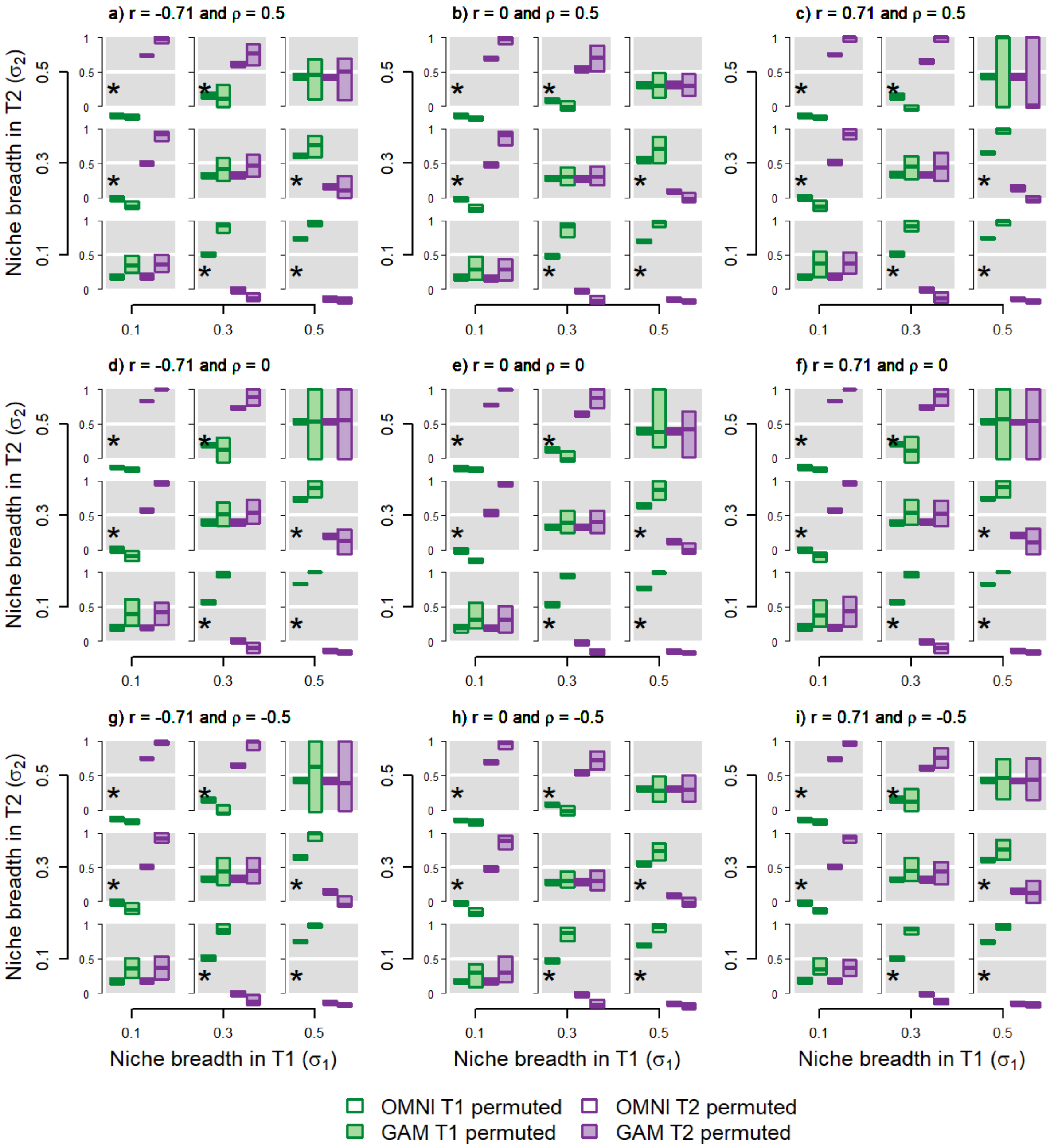


**Figure S7:** ***Experiments 7-9—Two influential variables modeled with GAMs***. Effect of niche breadth, collinearity, and niche covariance on inferences of variable importance using GAMs measured using $COR_{pa}$. Two variables, T1 and T2, each influence the niche according to the species' niche breadth in each. Each subpanel represents a landscape occupied by a species with the given niche breadths ($\sigma_1$ and $\sigma_2$), correlation between predictors ($r$) and niche covariance ($\rho$). Panel e corresponds to Fig. 4a in the main text (changing niche breadth with no correlation between predictors, no niche covariance). Asterisks indicate reliable discrimination and Latin crosses successful calibration. See Box 1 for further details on interpreting each subpanel in a panel.

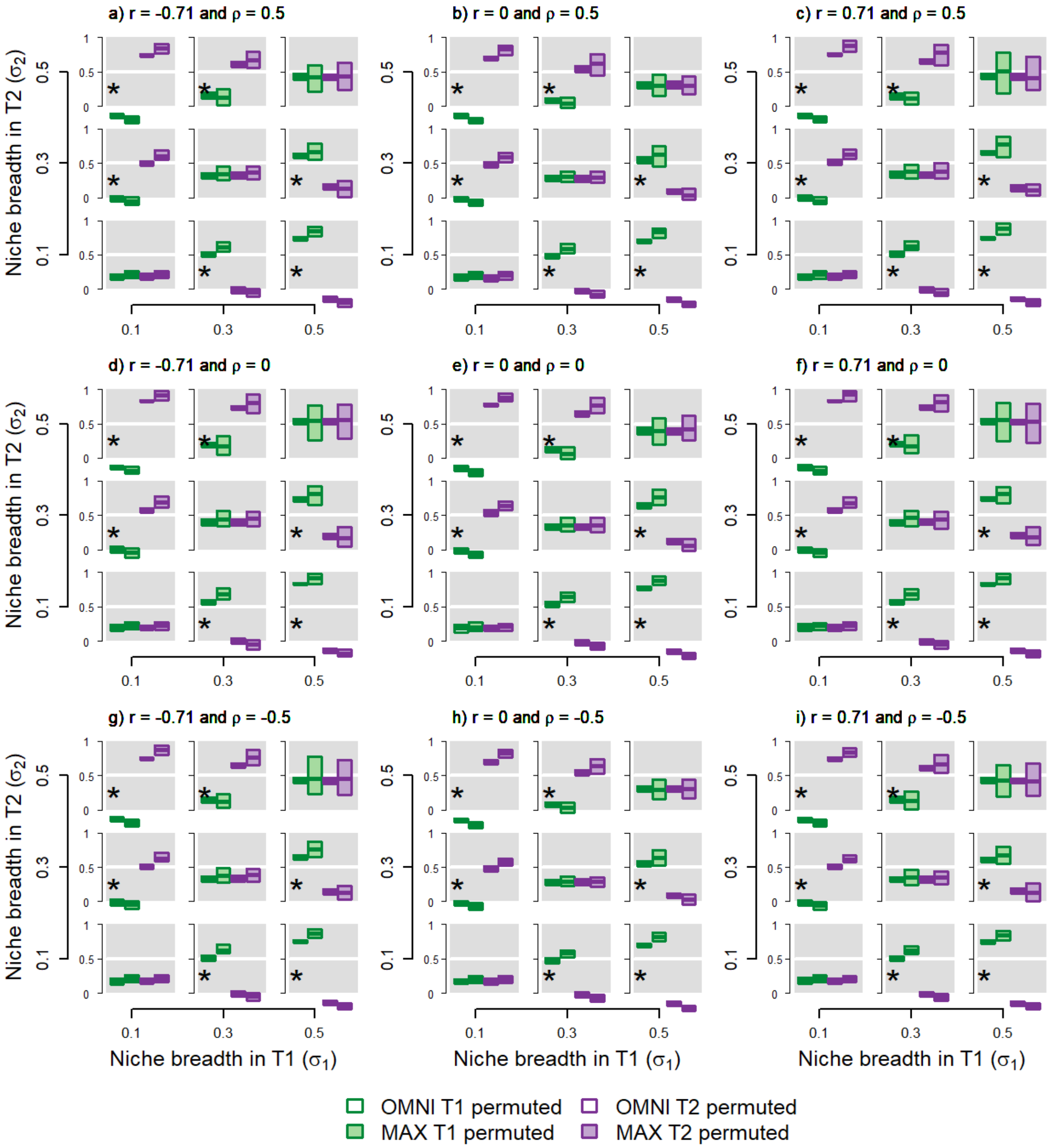


**Figure S8:** ***Experiments 7-9—Two influential variables modeled with Maxent***. Effect of niche breadth, collinearity, and niche covariance on inferences of variable importance using Maxent measured using $COR_{pa}$. Two variables, T1 and T2, each influence the niche according to the species' niche breadth in each. Each subpanel represents a landscape occupied by a species with the given niche breadths ($\sigma_1$ and $\sigma_2$), correlation between predictors ($r$) and niche covariance ($\rho$). Panel e corresponds to Fig. 4a in the main text (changing niche breadth with no correlation between predictors, no niche covariance). Asterisks indicate reliable discrimination and Latin crosses successful calibration. See Box 1 for further details on interpreting each subpanel in a panel.

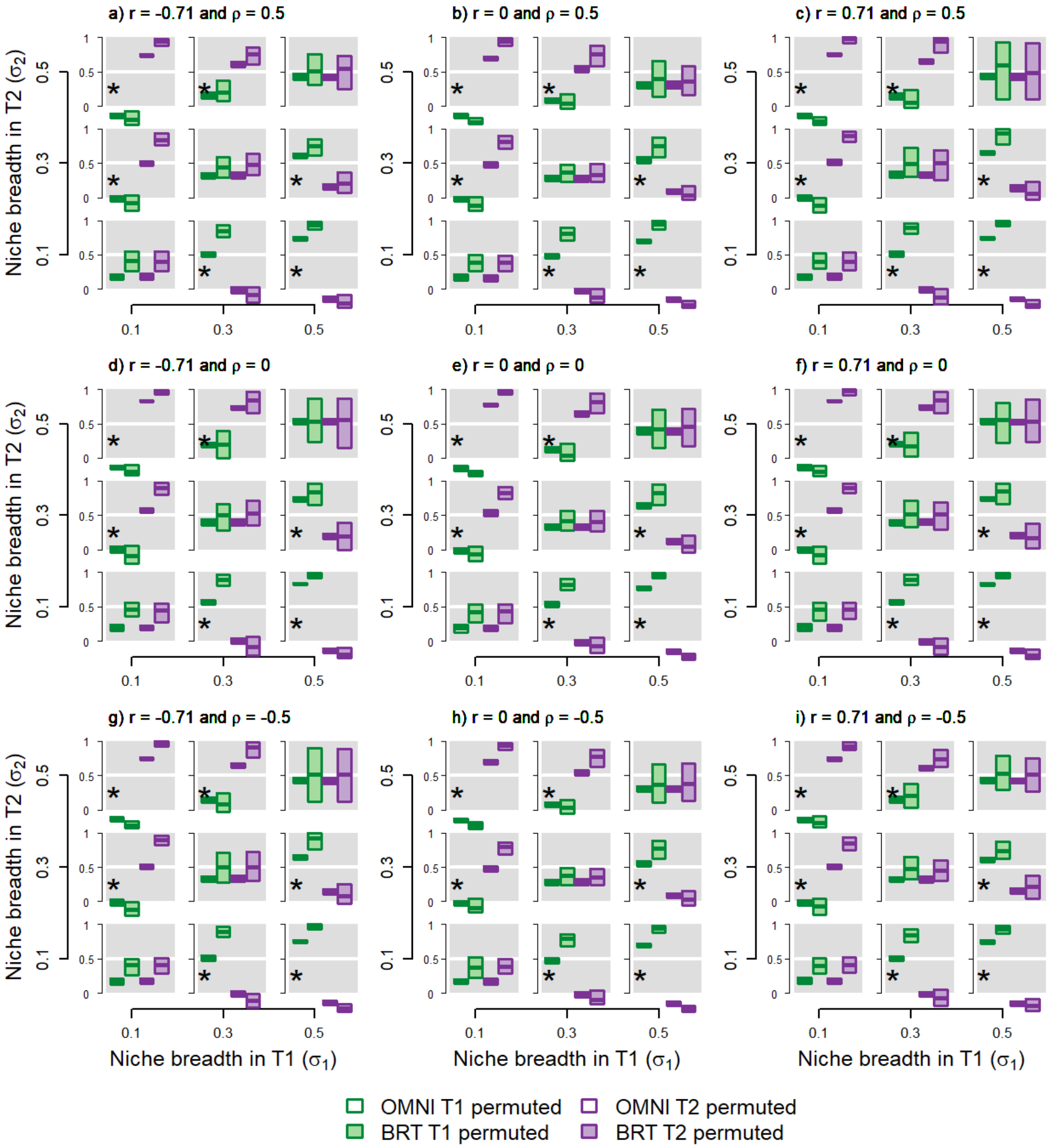


**Figure S9: *Experiments 7-9—Two influential variables modeled with BRTs*.** Effect of niche breadth, collinearity, and niche covariance on inferences of variable importance using BRTs measured using $COR_{pa}$. Two variables, T1 and T2, each influence the niche according to the species' niche breadth in each. Each subpanel represents a landscape occupied by a species with the given niche breadths ($\sigma_1$ and $\sigma_2$), correlation between predictors ($r$) and niche covariance ($\rho$). Panel e corresponds to Fig. 4a in the main text (changing niche breadth with no correlation between predictors, no niche covariance). Asterisks indicate reliable discrimination and Latin crosses successful calibration. See Box 1 for further details on interpreting each subpanel in a panel.

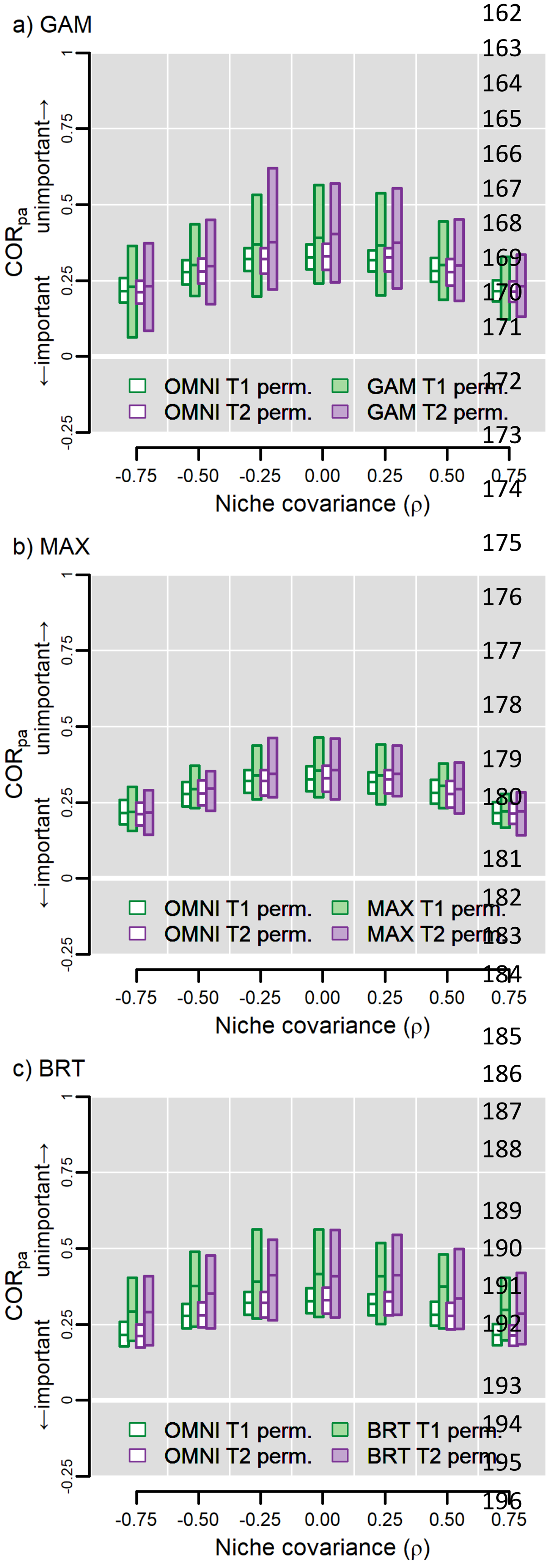


**Figure S10:** ***Experiment 8—Niche covariance.*** Variable importance increases as the magnitude of interaction between predictors increases. Here, niche width was held constant ($\sigma_1 = \sigma_2 = 0.3$) and the variables were uncorrelated on the landscape ($r = 0$). Since the variables had equal niche width, they should have the same influence. Latin crosses indicate successful calibration. See Box 1 for further details on interpreting each subpanel in a panel.
