## Appendix 9 for "Testing the ability of species distribution models to infer variable importance"

**Appendix 8: Algorithm-specific metrics of variable importance**

[Author names withheld for double-blind peer review.]

### Overview of methods

In the main text of the article we present results using the Continuous Boyce Index (CBI) as test statistic for the permute-after-calibration test of variable importance. In this supplement we present figures complementary to those in the main text except we show results using metrics specific to each algorithm. For BRTs, this is calculated by enumerating the frequency with which each variable appears across trees, weighting frequencies by the total reduction in deviance provided by the variable, and standardizing so the sum of all across all predictors equals 1 (Elith et al. 2008). Maxent has three algorithm-specific metrics. Two of these rely on "gain", which is the sum of predicted values at occurrences minus predicted values at background sites minus a regularization fitting penalty (Merow et al. 2013). Gain is unitless.

- *Gain*: This is the gain of the model with all variables included, without a focal variable, or with just the focal variable. In our case, we only examined scenarios with two variables, so gain of a model without a variable is the same as gain of a model with just the opposing variable.
- *Contribution importance*: Maxent finds an optimal solution by iteratively increasing the gain of the model through adding/subtracting a small amount from the coefficient for one term each iteration. This change in gain is cumulated for each variable that appears in the term. These values are then standardized across variables to sum to 1. Note that this metric is somewhat sensitive to the order in which variables are handled; a different programming instantiation that ends in the same final model might attribute different amounts of gain to predictors. Thus, the values are particular to the software used to calibrate the model.
- *Permutation importance*: This is the same as the permute-after-calibration test (Breiman 2001) using $AUC_{bg}$, except that training sites are used to calculate the test statistic, and values are standardized to sum to 1.

GAMs do not have an algorithm-specific metric of variable importance per se, but AIC-based model weights can be used to calculate variable-specific weights (Burnham & Anderson 2002; see also Murray & Conner 2009, Galipaud et al. 2014, and Giam & Olden 2016). We calibrated GAMs using an exhaustive evaluation of univariate and bivariate models (i.e., smooths for TRUE and FALSE individually or combined, plus univariate and bivariate smooths in the same model; described in Appendix 1). In almost all cases the model with the bivariate smooth (and no univariate smooths) had an AICc weight of ~1. Hence, both TRUE and FALSE appear to have equal weight using this method. As a result, we do not display algorithm-specific results for GAMs.

Results using the algorithm-specific metrics were qualitatively almost the same as using metrics based on predictive accuracy (CBI, AUC) or COR. However, in some cases responses were slightly different. For example, CBI, AUC, and COR all indicate that importance increases with the magnitude of niche covariance (Fig. 4b and Figs. S10 in Appendices 3-7. However, none of the algorithm-specific metrics displayed this kind of response (Fig. S8, this Appendix).

Note that neither BRT-specific nor Maxent-specific metrics can be calculated using OMNI (OMNI does not use training data so results using the permute-after-calibration test for OMNI are not strictly comparable to results using permutation of training data for Maxent).

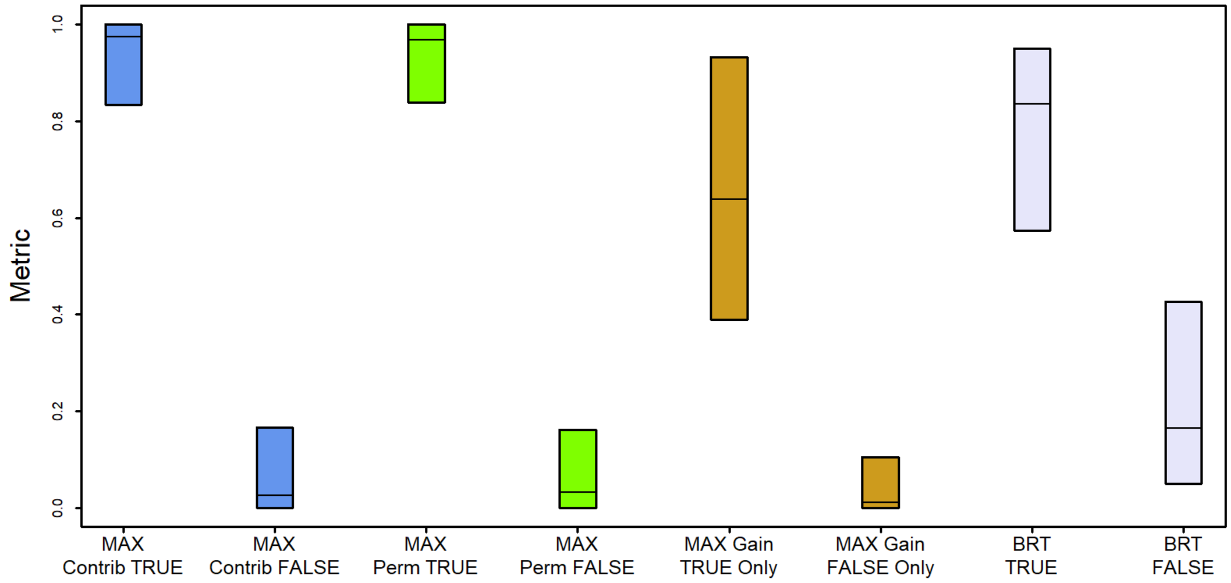


**Figure S1:** ***Experiment 1*****—Simple scenario**. Algorithm-specific tests of variable importance. All values except "gain TRUE only" and "gain FALSE only" have the range [0, 1]. Values for gain TRUE only" and "gain FALSE only" have been standardized to plot them on the same graph as the other metrics. Comparisons are only valid within a test (i.e., Maxent's "contribution" for TRUE versus FALSE). In each case TRUE is reliably discriminated from FALSE. Bars represent the inner 95% of test metric values. Horizontal lines in bars represent median values.

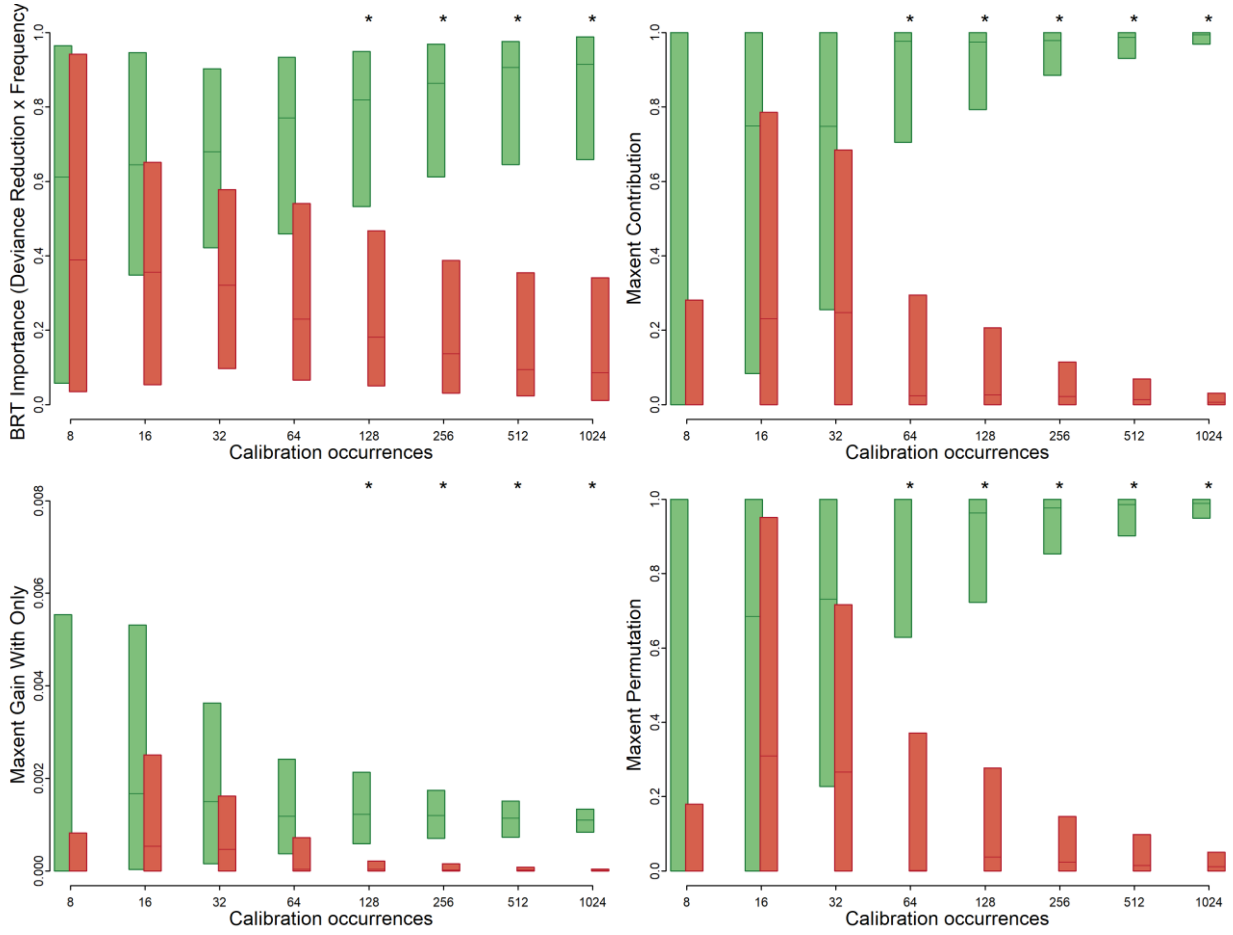


**Figure S2:** ***Experiment 2—Sample size***. Asterisks indicate successful discrimination between TRUE and FALSE. Green bars represent results for TRUE and red bars for FALSE. Bars represent the inner 95% of test metric values. Horizontal lines in bars represent median values.

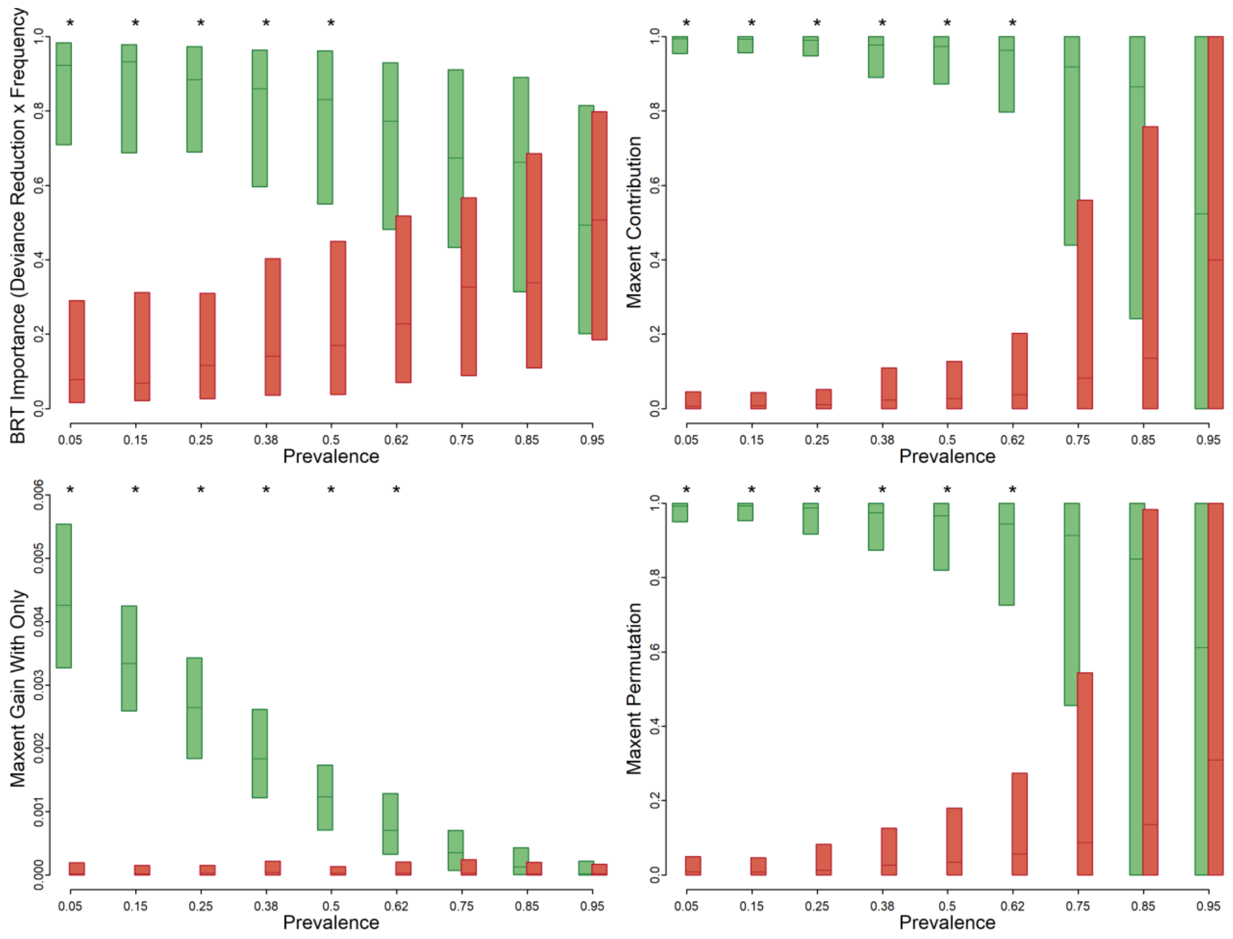


**Figure S3:** ***Experiment 3—Prevalence***. Asterisks indicate successful discrimination between TRUE and FALSE. Green bars represent results for TRUE and red bars for FALSE. Bars represent the inner 95% of test metric values. Horizontal lines in bars represent median values.

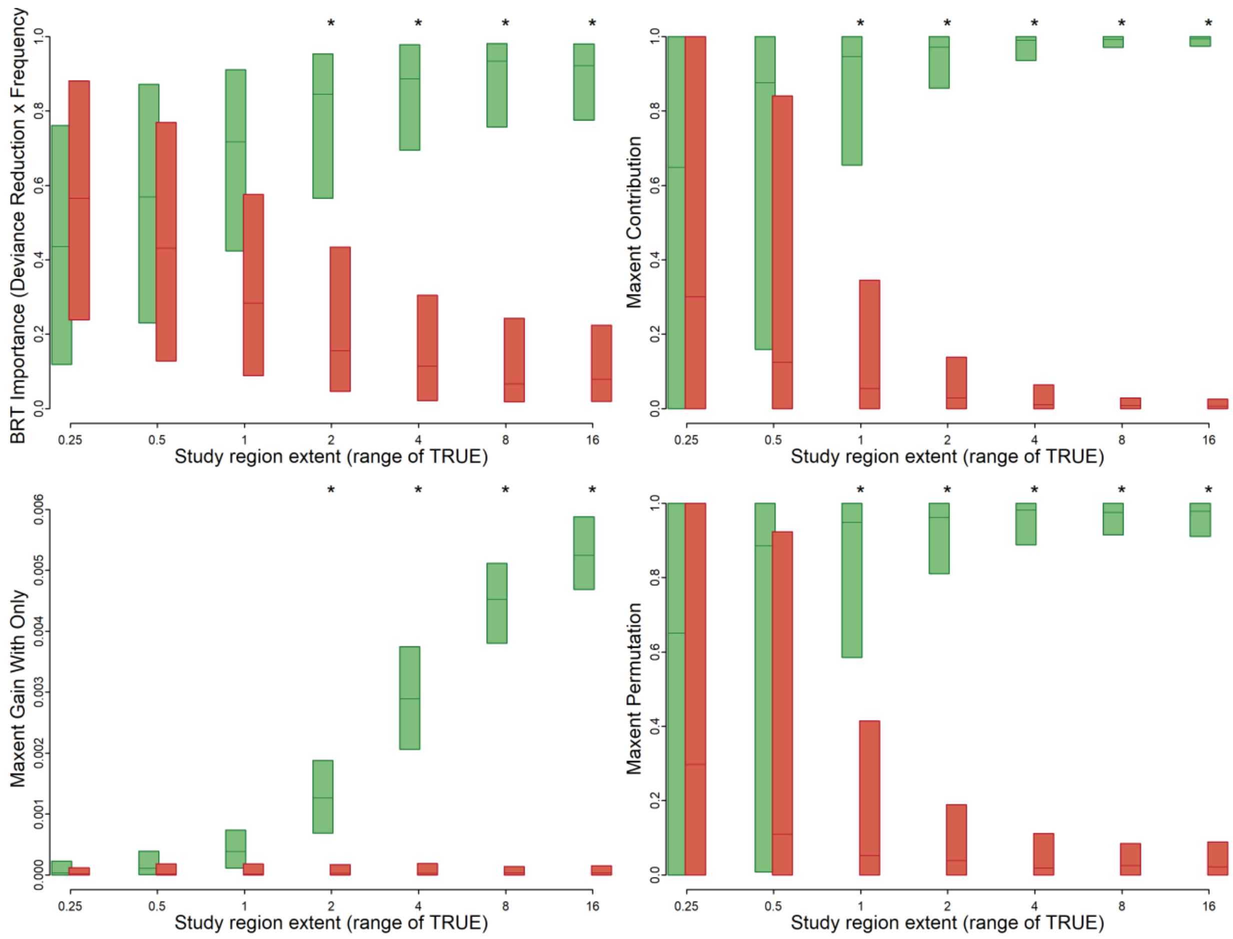


**Figure S4:** ***Experiment 4—Study region extent***. Asterisks indicate successful discrimination between TRUE and FALSE. Green bars represent results for TRUE and red bars for FALSE. Bars represent the inner 95% of test metric values. Horizontal lines in bars represent median values.

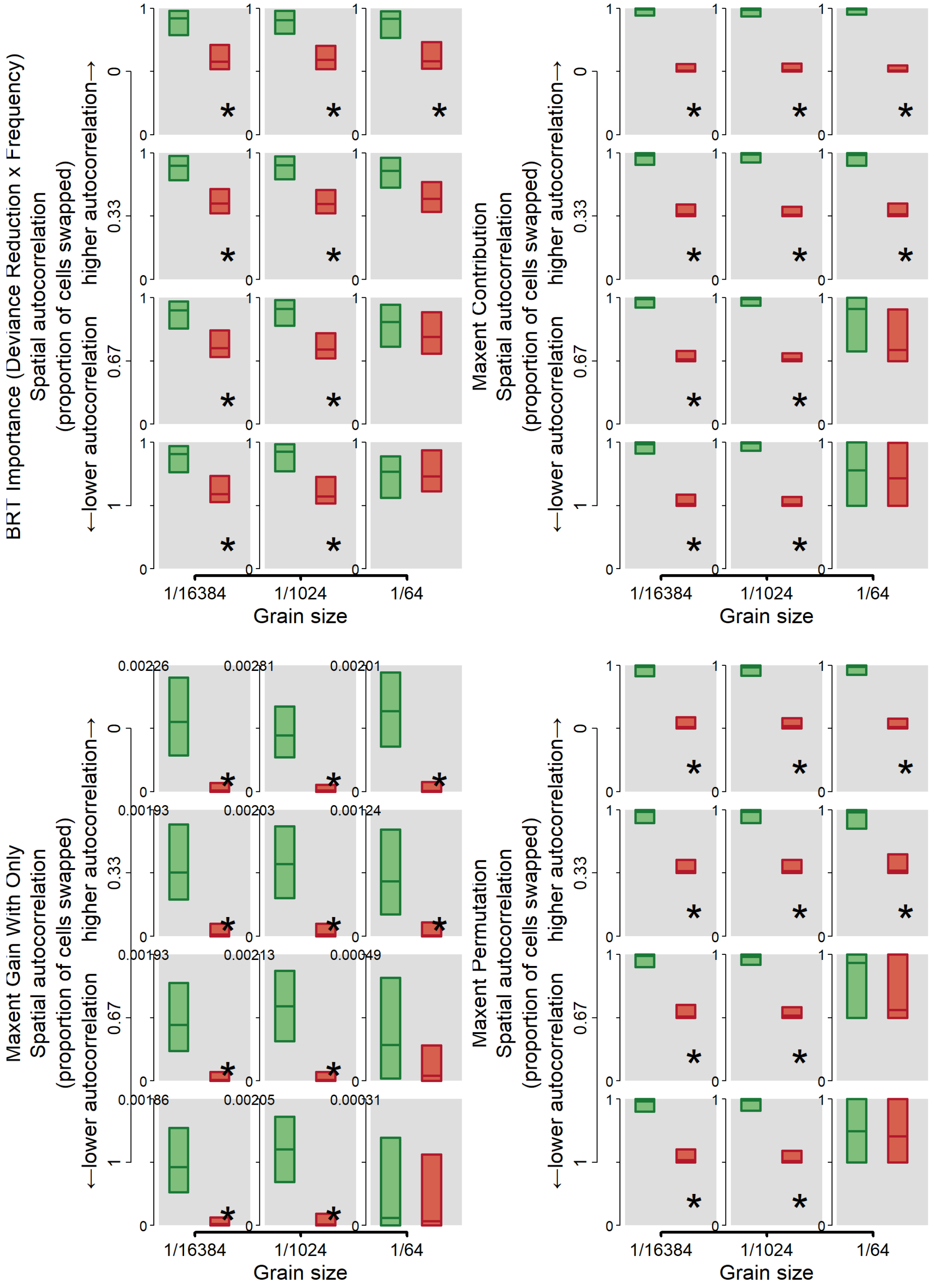


***Figure S5: Experiment 5—Spatial resolution and autocorrelation of environmental data*.** Asterisks indicate successful discrimination between TRUE and FALSE. Green bars represent results for TRUE and red bars for FALSE. Bars represent the inner 95% of test metric values. Horizontal lines in bars represent median values.

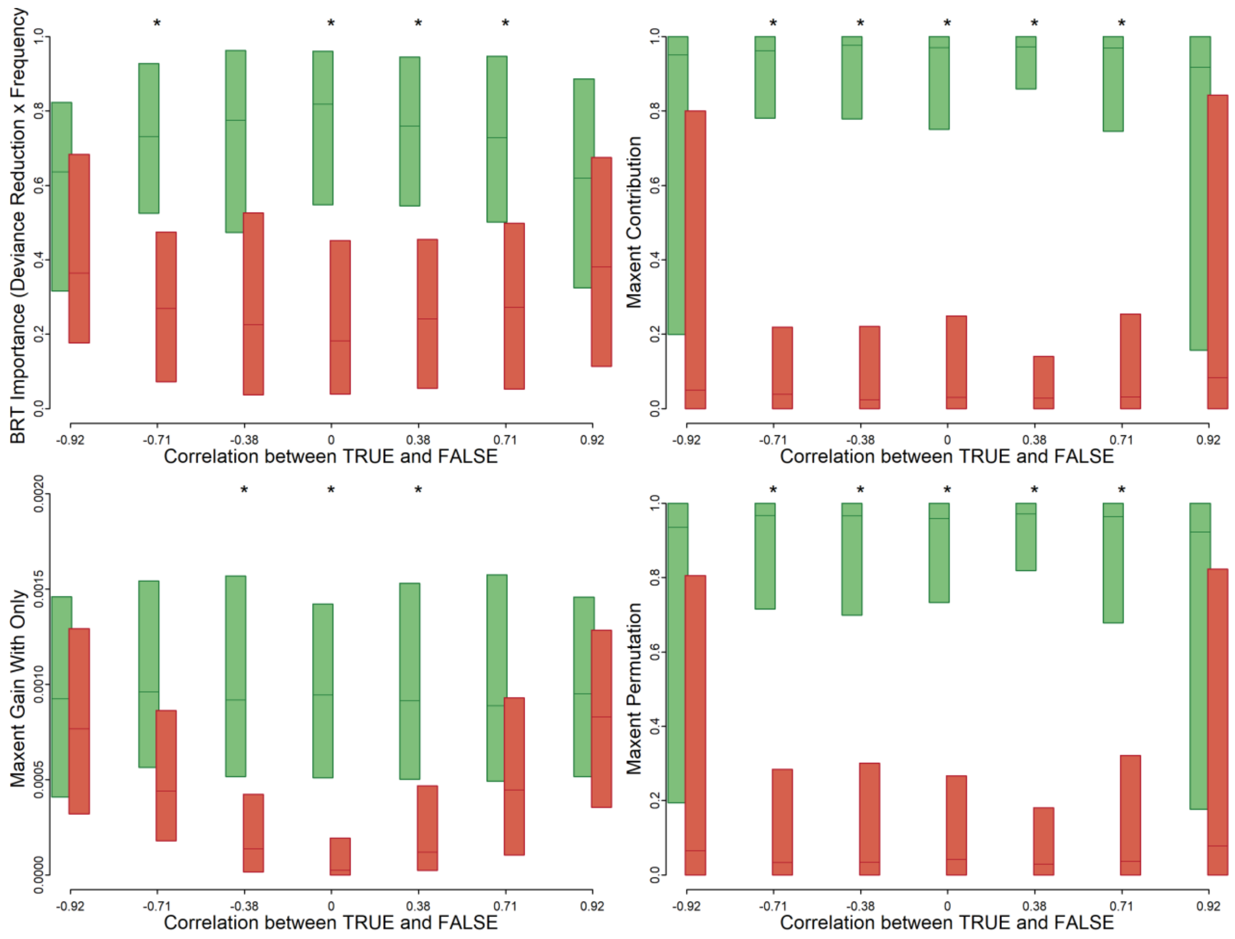


**Figure S6:** ***Experiment 6—Collinearity***. Asterisks indicate successful discrimination between TRUE and FALSE. Green bars represent results for TRUE and red bars for FALSE. Bars represent the inner 95% of test metric values. Horizontal lines in bars represent median values.

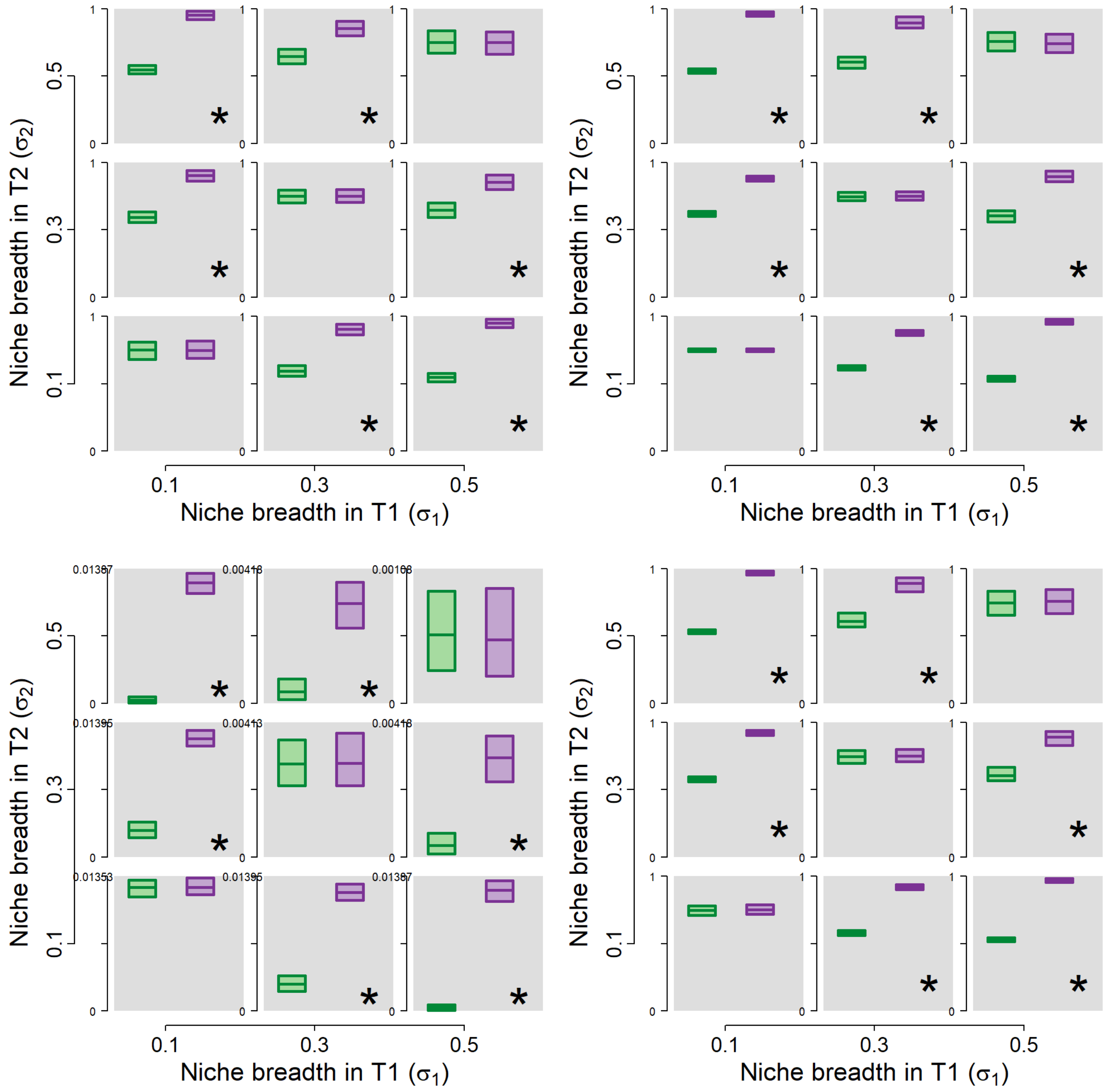


**Figure S7:** ***Experiment 7—Niche breadth.*** Asterisks indicate successful discrimination between TRUE and FALSE. Green bars represent results for TRUE and red bars for FALSE. Bars represent the inner 95% of test metric values. Horizontal lines in bars represent median values.

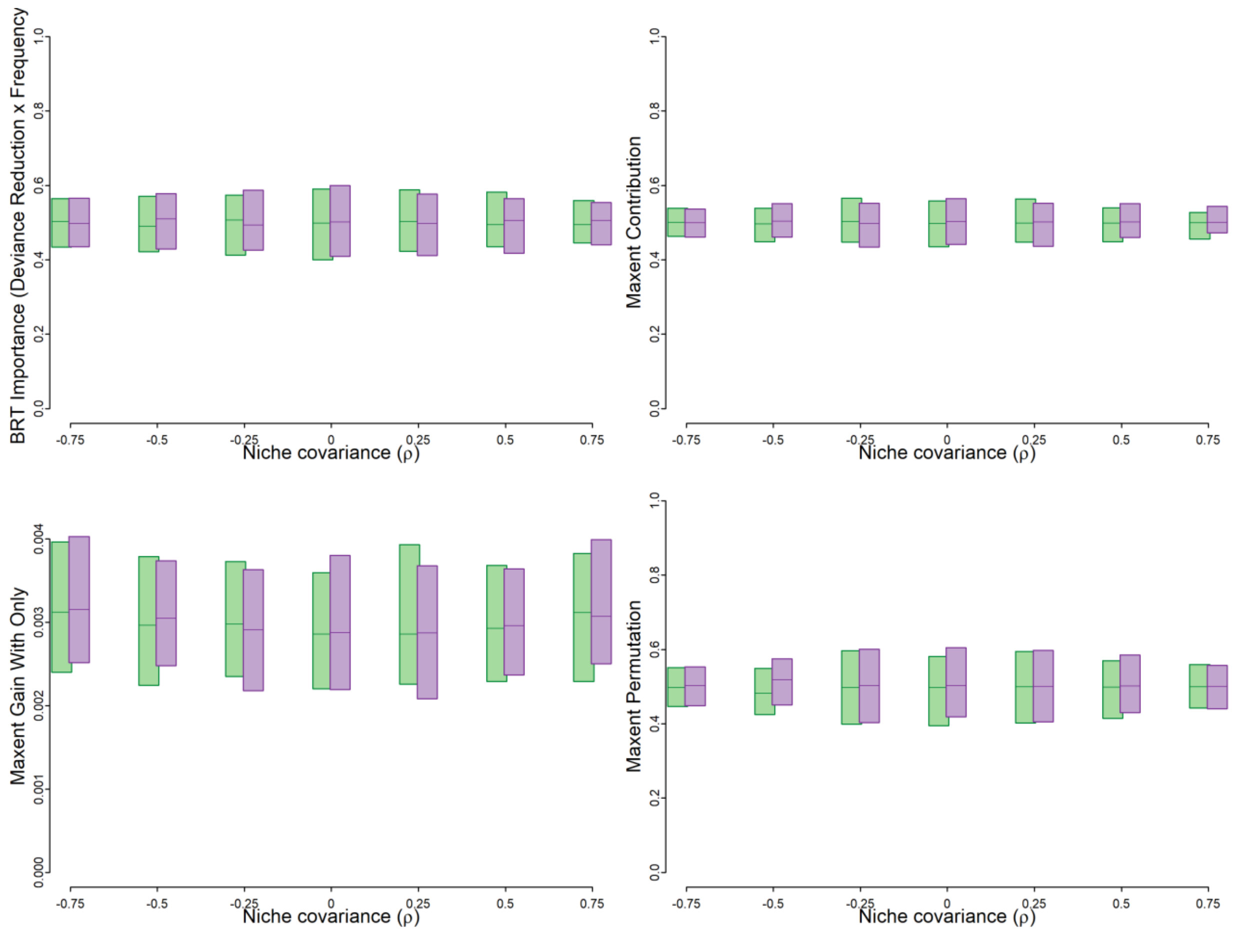


**Figure S8:** ***Experiment 8—Niche covariance***. Asterisks indicate successful discrimination between TRUE and FALSE. Green bars represent results for T1 and purple bars T2. Niche width was held constant ($\sigma_1 = \sigma_2 = 0.3$) and the variables were uncorrelated on the landscape ($r = 0$). Since the variables had equal niche width, they should have the same influence.
