## Appendix 10 for "Testing the ability of species distribution models to infer variable importance"

### Appendix 8: Sensitivity of the Continuous Boyce Index to improbable test presences

[Author names withheld for double-blind peer review.]

#### Overview of methods

In the main text of the article we focus on measures of variable importance using the Continuous Boyce Index (CBI; Boyce et al. 2002; Hirzel et al. 2006). CBI is a measure of a model's calibration accuracy (correlation of predictions with the actual probability of presence; Phillips & Elith 2010). There are some indications in the modeling literature that CBI may be a more sensitive measure of model performance than AUC (which is a measure of a model's discrimination accuracy (Cianfrani et al. 2010; Breiner et al. 2015), or ability to differentiate presences from absences or presences from background points; Phillips & Elith 2010). Here we focus on one aspect of sensitivity: the "leverage" of a single "improbable" test occurrence with a very low probability of occurrence on CBI. Since CBI is calculated using the Spearman rank correlation statistic (Boyce et al. 2002), we expect that including an improbable test site could dramatically reduce CBI even if there are many other presences with high predicted suitability.

To test our hypothesis, we constructed a simple situation in which 10,000 predictions at background sites were evenly spaced between 0 and 1. Predictions at "probable" test occurrence sites were also evenly distributed between 0.5 and 1. We simulated two cases. In the first case we used 200 probable presences as the test data. In the second case we used 199 probable presences plus a single improbable presence which was assigned a predicted suitability of 0.1, 0.01, 0.001, or 0.0001.

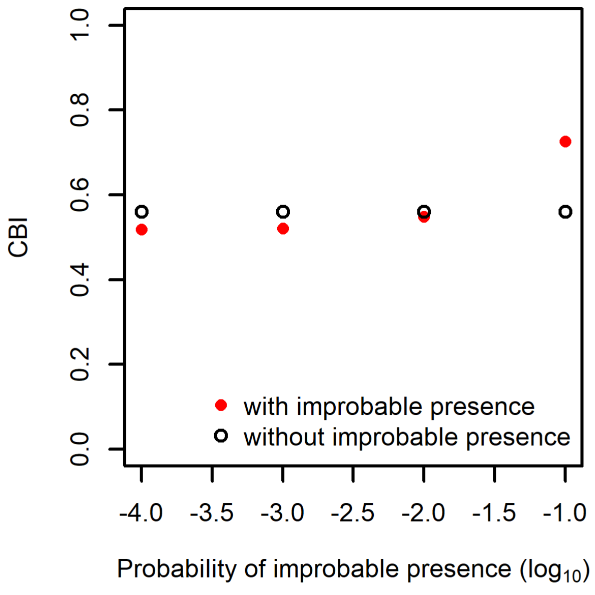


**Figure S1**. Including a single "improbable" presence in the test set can dramatically reduce CBI.

Including the single improbable presence from 0.1 to <0.1 reduced CBI dramatically (Fig. S1). Interestingly, including the single improbable presence with a probability of 0.1 elevated CBI above the

CBI calculated using just the probable presences. By default, the *contBoyce* function in the *[redacted for peer review]* package [redacted for peer review] drops bins without test presences (we used the default in evaluating models). Hence, the predicted-to-expected ratio used to calculate CBI is tabulated only in the bins with the improbable presence and the probable presences. As a result, adding an improbable presence increases the number of bins (sample size), which generally increases the Spearman rank correlation coefficient.

When a landscape is large (as when the extent of a study region is large; *Experiment 4: Effect of study region extent on assessing variable importance*), the low probability of presence in highly unsuitable areas can be offset by the size of these areas. In this case improbable presences become more likely. When this is the case and the improbable presence has a very low predicted probability of occurrence, then its inclusion can reduce CBI disproportionately. As a result, CBI may actually be *too* sensitive to outliers to serve as a robust indicator of model performance in all cases.

[redacted for peer review]
